# Head dopaminergic and serotonergic neurons monitor alterations in gut bacterial activity through a gut-brain-gut axis

**DOI:** 10.1101/2024.06.17.599455

**Authors:** Guanqun Li, Yangyang Wu, Xiaowen Huang, Minghui Du, Hongyun Tang

## Abstract

The brain is suggested to monitor gut microbial changes to critically maintain health, whereas the particular bacterial alterations and the mechanisms involved are largely unclear. Here, through a genome-wide screen, we identified 29 *E. coli* genes whose inactivation can be surveilled by *C. elegans* head dopaminergic and serotonergic neurons through increasing host neuronal dopamine and serotonin biosynthesis. Mechanistically, we found that these head neurons respond to the *E. coli* lack of respiratory chain genes by detecting the bacteria-caused reductions in host labile iron levels that impair intestinal mitochondria, and the intestine-expressed mitochondrial stress sensor ATFS-1 is critically involved. Furthermore, this neuronal response in turn promotes intestinal ferritin-1 expression to counteract bacteria-caused labile iron reduction and thereby maintaining mitochondrial function. Our findings systematically identify bacterial activity changes that elevate brain dopamine and serotonin levels and unveil an unexpected gut-brain-gut axis in which the head neurons inspect bacterial activity-mediated changes in host iron metabolism and mitochondrial function, enlightening crucial roles and mechanisms of brain in surveilling microbial activity alterations.

## Introduction

Throughout the life of an animal, the activity and community composition of gut microbes change in numerous ways, which is implicated to profoundly affect the host physiological activities ^1–4^. To maintain the health of host, the brain has been suggested to monitor these changes in gut bacteria and orchestrate appropriate responses ^1,5–7^. Accumulating evidence, including the results from fecal microbes transplantation experiments, supports that host nervous system respond to the changes in the gut microbial composition to adjust various physiological processes, which is important for promoting health such as relieving behavioral abnormalities and maintaining gut homeostasis ^1,8–11^. However, whether and how the alterations in gut microbial activity, occurring in response to various host and environmental factors, such as aging, diet and drug treatment ^1–3^ and thereby contributing heavily to the diversity of bacterial stimulations that act on the host, can be monitored by the nervous system has barely been explored. For example, metabolic activity of gut bacteria exerts influence on host through various means such as facilitating the digestion and metabolism of the host and generating effective metabolites ^12–14^ and thus is speculated to critically impact host biological processes, but a full investigation of the potential of these microbial metabolism changes in altering activity of the nervous system is lacking. Therefore, systematically examining microbial activity change-mediated neuronal responses and elucidating the underlying mechanisms are of great importance, which should provide comprehensive insights into understanding the microbes-brain interactions.

Remarkably, gut bacteria are implicated to be able to alter biosynthesis of neurotransmitters to profoundly affect host physiology ^1,15,16^. For example, it was reported that gut microbiota promotes serotonin production from the colonic enterochromaffin cells to affect the gastrointestinal motility in mouse ^15^. However, whether the gut bacteria could increase biosynthesis of neurotransmitters such as serotonin and dopamine in the brain, which in turn may modulate host physiological functions to promote health, remains largely elusive ^1,16^. Especially, the particular bacterial changes that could promote the levels of the brain neurotransmitters and the underlying mechanisms are unclear. Identifying bacterial activity changes that are capable of increasing brain dopamine and serotonin biosynthesis is highly important, which not only sheds light on understanding how gut microbes improve neurologic functions but also holds promise to develop strategies to ameliorate the defects caused by decrease in dopamine and serotonin.

The communications between microbes and the nervous system are very complex, in which the gut bacteria interact with the nervous system through nervous, endocrine, and immune communications with the critical involvement of interorgan crosstalk ^1,17^. Emerging evidence supports that gut, the major organ that directly interacts with bacteria, likely plays a dominant role in mediating the perceiving of the gut bacteria by the host nervous system, which could orchestrate feedback responses to modulate the gut physiology and its associated microbes ^5,10,18^. Considerable researches support that when confronting bacterial changes, the gut may generate various immunological and metabolic factors that ultimately act on the neurons and thereby influencing the brain activities ^1,18,19^, whereas the exact mechanisms are yet to be fully understood. For instance, it is still understudied what intestinal alterations need to be conveyed to the nervous system when confronting bacterial change, which is critical for understanding the implicated physiological importance of this interaction. Moreover, the communications between gut bacteria and the brain are bidirectional ^1^, but the underlying gut-brain-gut regulatory loop for any specific physiological functions are largely unclear. Altogether, due to the tremendous diversity of bacterial stimuli as well as the complex connections between bacteria and the host nervous system, in-depth research is imperative to illustrate the cellular and molecular pathways underlying bacteria-neuron communications and the relevant physiological functions.

*C. elegans* is widely used to study the basic rules underlying the communications between bacteria and the host ^6,20–22^, which has led to seminal discoveries, such as bacteria-mediated life-span extension and microbes-induced change of host behavior ^6,23^, enlightening the mechanisms as well as the important physiological roles of microbes-host interactions. Here, by using *C. elegans* as a model system, we aimed to obtain an overall understanding of what alterations in bacterial activities can trigger elevation of dopamine and serotonin biosynthesis in the head neurons and to identify the underlying molecular and cellular mechanisms. As mutations in bacterial genes can result in changes in microbial activities, we administered single-deletion *E. coli* mutants from the Keio library containing knockouts of all non-essential bacterial genes to *C. elegans* and assessed responses from the head dopaminergic and serotonergic neurons by analyzing the induction of BAS-1, a shared enzyme of dopamine and serotonin biosynthesis. We found that deletions of bacterial genes involved in a variety of processes, including those encoding respiratory chain components, triggered response from these head neurons, as measured both by increased fluorescence of a GFP-fused BAS-1 transgenic reporter and by the elevation of dopamine and serotonin level. Furthermore, we discovered that the head dopaminergic and serotonergic neurons perceived the presence of the bacteria containing mutations in respiratory chain genes by inspecting the resultant reduction in peripheral labile iron levels. Mechanistically, we found that there exists a gut-brain-gut axis which allows the head neurons to detect bacteria-induced decreases in labile iron levels and the consequent impairment of the intestinal mitochondria and then to orchestrate appropriate responses to prevent further decreases in available iron levels in the gut, which is important for maintaining mitochondrial function of the host. Overall, our study systematically identified the changes in bacterial activity that provoke head dopaminergic and serotonergic neuronal response to increase dopamine and serotonin biosynthesis and uncovered an unexpected gut-brain-gut axis playing a key role in monitoring bacterial activity alterations, which exemplifies a bidirectional communication mechanism by which the brain monitors changes in the activities of the gut and gut-associated bacteria.

## Results

### Head serotonergic and dopaminergic neurons in *C. elegans* respond to changes in gut bacterial activity, including the inactivation of *E. coli* respiratory chain genes, by increasing neuronal dopamine and serotonin biosynthesis

Throughout an animal’s life, the activity of gut bacteria can change in numerous ways to potentially affect host physiology, and the nervous system may monitor these alterations in microbial activity to maintain health. To systematically investigate what types of changes in microbial activities could provoke responses from the host nervous system, we carried out a screen to explore whether treating *C. elegans* with any of the 3985 *E. coli* mutants containing single deletions of all the nonessential genes from the Keio collection ^24^ could trigger responses in two sets of neurons in the head of *C. elegans,* the serotonergic and dopaminergic neurons, to increase the biosynthesis of their corresponding neurotransmitters. In this screen, we assessed the expression of BAS-1, the shared enzyme for both dopamine and serotonin biosynthesis that is expressed in these neurons ^25^ (Fig. 1A), by measuring the fluorescence intensity of GFP-fused BAS-1 driven by the native *bas-1* promoter from a previously-used *Is[bas-1p::BAS-1::GFP]* transgene (Fig. 1B) ^26^. After three rounds of testing, single deletions of 29 bacterial genes were found to lead to higher BAS-1 expression in the *C. elegans* head neurons than in the control worms treated with the wild-type parent *E. coli* strain BW25113 (Fig. 1B; Supplementary information, Fig. S1A). The identified microbial genes encode a variety of classes of proteins, including transporters, enzymes and membrane proteins, and GO analyses indicated that these bacterial genes were mainly involved in metabolic processes and transport, including the electron transport chain (Supplementary information, Fig. S1 B-D; Fig. 1C). These findings indicate that the inactivation of bacterial genes involved in various biological processes triggers responses from host dopaminergic and serotonergic neurons.

**Fig. 1:**
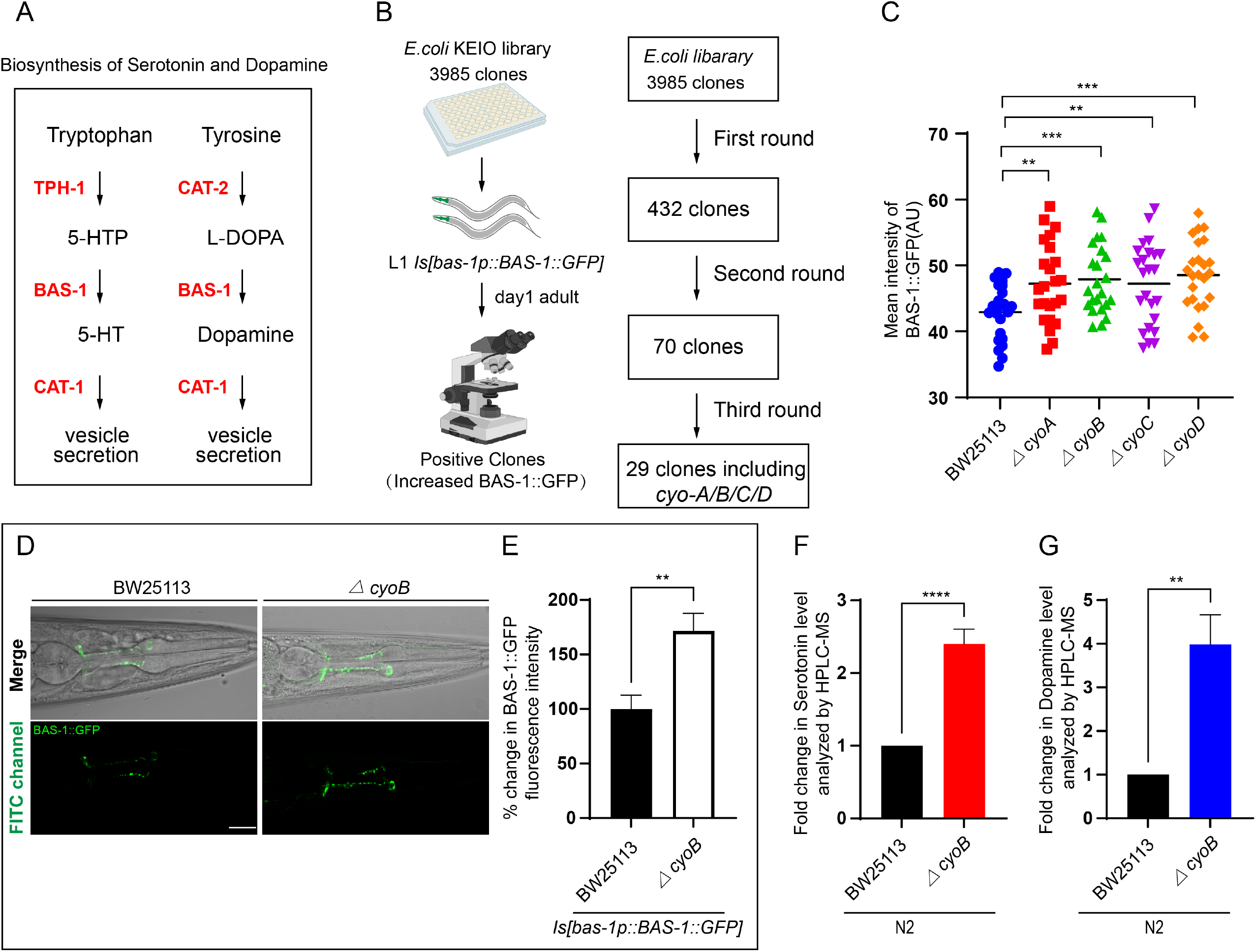
Head serotonergic and dopaminergic neurons of *C. elegans* respond to bacterial activity changes, including the inactivation of *E. coli* respiratory chain genes, to increase biosynthesis of serotonin and dopamine. **(A)** Schematic diagram indicating the biosynthesis process of serotonin and dopamine neurotransmitters. 5-HTP, 5-HT and L-DOPA indicate 5-hydroxyl-L-tryptophan, serotonin and L-dihydroxyphenylanaline, respectively. TPH-1 exhibits tryptophan 5-monooxygenase activity. CAT-2 enables tyrosine 3-monooxygenase activity. BAS-1, enable L-dopa decarboxylase activity, is the shared enzyme for both dopamine and serotonin biosynthesis. CAT-1 transports dopamine and serotonin into synaptic vesicles. **(B)** Illustration of the genome-wide screen of the bacterial genes that enhanced *bas-1p::BAS-1::GFP* expression in the host *C. elegans* upon inactivation. The Keio library containing 3985 single deletions of all the nonessential genes of *E. coli* was used to screen for bacterial mutants that enhanced the expression of BAS-1, the shared enzyme for both dopamine and serotonin biosynthesis, as indicated in (A). The bacterial clones that significantly enhanced the expression of BAS-1::GFP in all three rounds of testing were defined as positive hits. **(C)** Quantitative analyses of BAS-1::GFP expression induced by feeding *Is[bas-1p::BAS-1::GFP] C. elegans* bacteria with a single deletion of *cyoA/B/C/D.* Wild-type BW25113 *E. coli*, the parental strain for generating these mutations ^24^, was used as a negative control for analyzing the effect of these genes deletion on BAS-1::GFP expression in the *C. elegans* host. Each dots represent the mean intensity of BAS-1::GFP expression in each worm. n > 20 for each group. **(D and E)** Images and bar graph indicating that *E. coli* with *cyoB* deletion (*ΔcyoB)* causes increased expression of BAS-1::GFP in *C. elegans* dopaminergic and serotonergic neurons. Wild-type worms carrying *Is[bas-1p::BAS-1::GFP]* transgene were analyzed. The bar graph shows quantification of the BAS-1::GFP fluorescence intensity which is relative to wild-type animals treated with BW25113 presented in (D). n > 30 for each group. Scale bar, 20 μm. **(F and G)** Bar graph showing HPLC-MS analyses of serotonin and dopamine levels in day-1 adult N2 *C. elegans* treated with BW25113 or *ΔcyoB E. coli*. The fold change of serotonin (F) or dopamine (G) was calculated by normalizing to the levels from the *C. elegans* treated with BW25113. n > 2000 each group. For all the panels, experiments were performed for at least three times and data are the mean ± SEM. **p < 0.01, ***p < 0.001 and ****p < 0.0001 by unpaired t test.

Interestingly, the bacterial mutants obtained from the screen were obviously enriched in bacterial genes encoding the cytochrome bo (3) ubiquinol oxidase subunit involved in the *E. coli* respiratory chain pathway, including *cyo-A/-B/-C/-D* (Fig. 1B and 1C). As an example bacterial mutant that strongly induced BAS-1::GFP, the *E. coli cyoB* deletion mutant (*ΔcyoB E. coli*) was used to further understand the detailed mechanism by which the head dopaminergic and serotonergic neurons respond to the bacterial activity changes (Fig. 1C). First, our further independent experiments corroborated the obvious induction of BAS-1::GFP by the *ΔcyoB E. coli* (Fig. 1D and 1E). Next, we assess whether this increase of BAS-1 level could promote corresponding neurotransmitters biosynthesis by HPLC-MS analyses. Indeed, we found that the serotonin and dopamine levels in the wildtype worms treated with *ΔcyoB E. coli* were approximately 2-fold and 4-fold of the levels in the animals cultured with BW25113 control, respectively (Fig. 1F and 1G). Overall, these results demonstrate that head serotonergic and dopaminergic neurons in *C. elegans* elevate biosynthesis of corresponding neurotransmitters in response to alterations in various microbial activities, including the inhibition of *E. coli* respiratory chain genes.

### Head serotonergic and dopaminergic neurons perceive the inactivation of the bacterial respiratory *cyoB* gene by detecting bacteria-caused decrease in host labile iron level

We then aimed to understand how the dopaminergic and serotonergic neurons could perceive the presence of *ΔcyoB E. coli* to upregulate BAS-1 expression. It has been suggested that neurons could monitor the dynamics of the microbiota by detecting the bacteria themselves or by detecting bacteria-caused host changes. Since competition for iron occurs commonly between bacteria and the host ^22,27^, we first tested whether bacteria might cause a change in iron metabolism to trigger the neuronal expression of *bas-1* in the host. To this end, we measured the cellular labile iron level, which provides available iron for metabolic demands ^28^, by the widely-used Calcein-AM staining method. When entering the cell, Calcein-AM forms cell-impermeable fluorescent Calcein that is quenched by labile iron; thus, high Calcein fluorescence indicates relatively low labile iron levels ^21,28^. 2-2’ bipyridyl (BP), an iron chelator commonly used to reduce the labile iron pool in *C. elegans* ^29^, was used as positive control, and compared to the N2 wild type worms treated with wild type bacteria and solvent, the ones treated with wild type bacteria and BP indeed displayed an increase in Calcein fluorescence (Fig. 2A a-d and 2B). Interestingly, an increase in Calcein fluorescence was observed in the *ΔcyoB E. coli-*treated wild type N2 *C. elegans* when compared to the BW25113-treated ones (Fig. 2A e-h and 2B), which indicates that *ΔcyoB E. coli* causes a decrease in intracellular labile iron level in these worms. Then, we confirmed this labile iron reducing effect of the *ΔcyoB E. coli* by showing that elevating intracellular labile iron levels by iron supplementation (FeCl_3_) was able to reverse the increase of Calcein fluorescence in the *ΔcyoB E. coli-*treated wild type N2 *C. elegans* to the level observed in the N2 worms treated with wildtype bacteria (Fig. 2A i-l and 2B). Furthermore, iron availability analyses were also performed by analyzing transcription level of *smf-3*, a homologue to the human divalent iron transporter DMT-1, which was known to be transcriptionally activated during iron deficiency ^30^. We found that, compared to BW25113-treated worms, *ΔcyoB E. coli-*treated N2 worms displayed increased *smf-3* mRNA level (Supplementary information, Fig. S2A), in agreement with the observation that *ΔcyoB E. coli* caused decrease in the levels of available iron. This induction of *smf-3* transcription by *ΔcyoB E. coli* could be totally recovered by iron supplementation (Supplementary information, Fig. S2A), further demonstrating the labile iron reducing effect of this bacterial strain. Altogether, these data suggest that *ΔcyoB E. coli* causes a reduction in intracellular labile iron levels in *C. elegans* host.

**Fig. 2:**
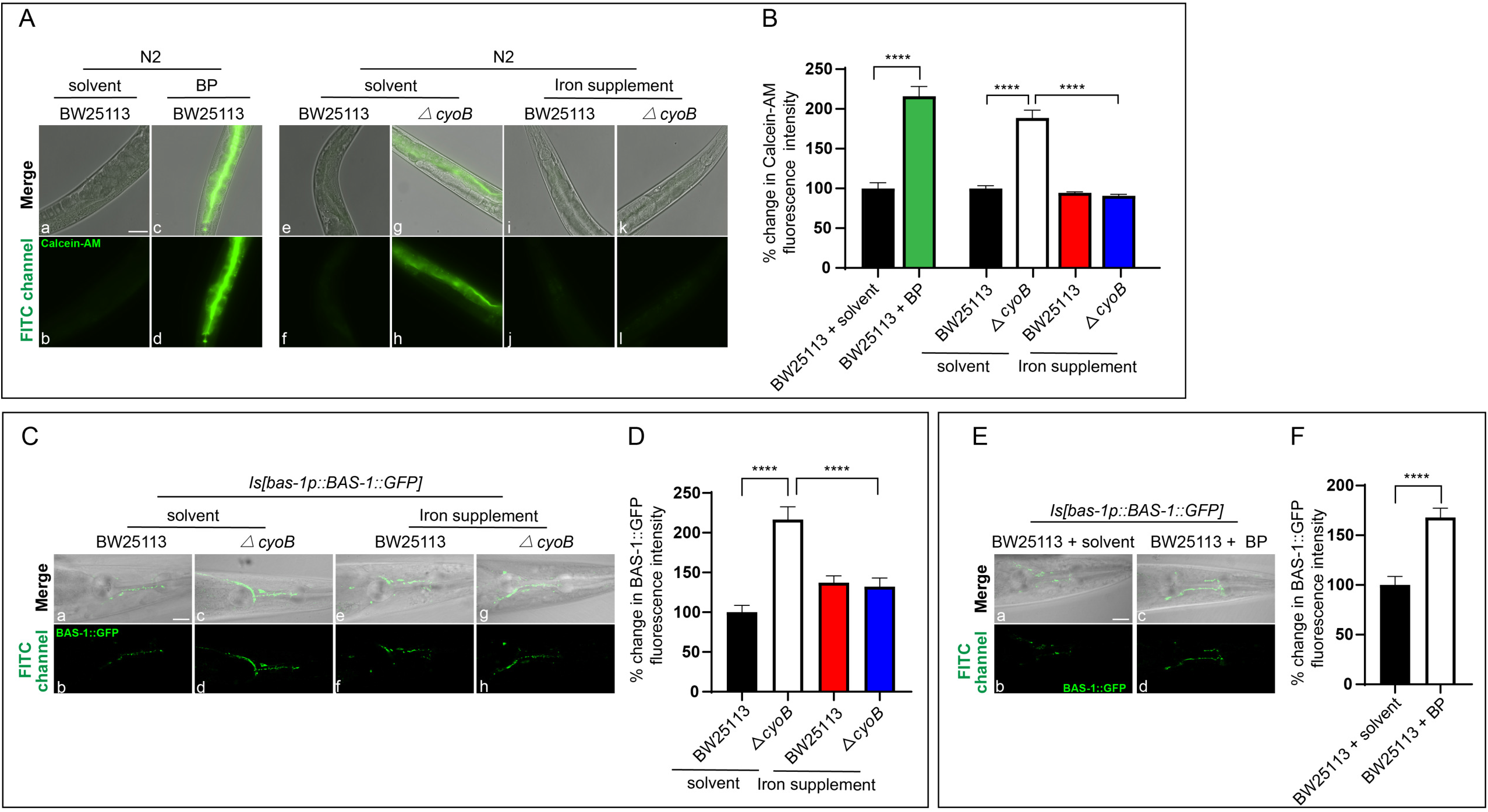
Serotonergic and dopaminergic neurons respond to inactivation of the bacterial respiratory chain gene *cyoB* by detecting bacteria-caused host labile iron insufficiency. **(A and B)** Images and bar graphs indicating the decrease in labile iron levels in the host *C. elegans* treated with *the ΔcyoB E. coli*. The intracellular labile iron level was measured by the widely used dye Calcein-AM, which enters cells and is immediately cleaved by intracellular esterases to form fluorescent Calcein, which is quenched by iron binding; thus, the increased fluorescence indicates a relatively low labile iron level. The iron chelator 2-2’ bipyridyl (BP) caused an increase in Calcein fluorescence (c, d). The increase in Calcein fluorescence induced by *the ΔcyoB E. coli* mutant (g, h) was reversed by supplementation with iron (4mM FeCl_3_) (k, l). The bar graph shows percent changes in the Calcein fluorescence intensity normalized to the levels in animals treated with both BW25113 and solvent. ****p < 0.0001 by one-way ANOVA and unpaired t test; n > 30 for each group. Scale bar, 50 μm. **(C and D)** Images and bar graphs showing that iron supplementation reverses the induction of BAS-1::GFP expression caused by *ΔcyoB* bacteria. The percent change of BAS-1::GFP fluorescence intensity was calculated by normalization to the levels in the animals treated with both BW25113 and solvent. Data are the mean ± SEM. ****p < 0.0001 by one-way ANOVA. n > 20 for each group. Scale bar, 20 μm. **(E and F)** Images and bar graph depicting the induction of BAS-1::GFP expression by iron-chelator 2-2’ bipyridyl (BP). BP was used to sequester the labile iron. The bar graph shows the % change in BAS-1::GFP fluorescence intensity calculated relative to the levels in the solvent-treated animals. Data are the mean ± SEM. ****p < 0.0001 by unpaired t test. n > 20 each group. Scale bar, 20 μm.

Next, we explored whether the observed increase in BAS-1 expression in the head neurons of worms treated with *ΔcyoB E. coli* was in response to the resultant reduction in labile iron levels. Indeed, when growing on the plates supplemented with iron, the BAS-1::GFP level from the *ΔcyoB E. coli*-treated *Is[bas-1p::BAS-1::GFP]* transgenic worms was similar to that observed in the ones treated with wildtype bacteria (Fig. 2C and 2D), indicating that elevating the labile iron level by this iron supplementation was able to almost completely suppress the *ΔcyoB E. coli*-induced increase in BAS-1 expression. Furthermore, we investigated whether reducing labile iron levels in *C. elegans* by iron chelator was able to induce *bas-1* expression in the dopaminergic and serotonergic neurons. Indeed, in the *Is[bas-1p::BAS-1::GFP]* worms cultured with wild-type BW25113 *E. coli,* the expression of BAS-1::GFP was higher in the BP-treated ones than in the solvent-treated ones (Fig. 2E and 2F), as that observed in the *ΔcyoB E. coli-*treated worms (Fig. 1D and 1E). Therefore, these results indicate that labile iron insufficiency triggers a response in *bas-1-*expressing neurons. Overall, these findings indicate that the inactivation of the microbial gene *cyoB* induces a decrease in labile iron levels in the host, which is detected by dopaminergic and serotonergic neurons to upregulate *bas-1* expression.

### The increase in neuronal dopamine and serotonin biosynthesis is crucial for the *C. elegans* host to counteract the bacteria-induced reduction in labile iron levels

Having shown that serotonergic and dopaminergic neurons respond to decreased labile iron levels, we next sought to determine the function of the upregulation of BAS-1 in these neurons in response to *ΔcyoB E. coli* and to assess which neuron type mediated this response. One possibility is that these head neurons monitor bacteria-caused reductions in iron levels to orchestrate a proper response to maintain iron homeostasis. Thus, we tested whether inhibited *bas-1* expression could exacerbate the reduction in labile iron levels in *ΔcyoB E. coli*-treated worms. As predicted, in the presence of *ΔcyoB E. coli,* the worms containing a loss-of-function (lf) mutation in *bas-1* displayed a further reduction in labile iron levels as they exhibited a dramatically higher Calcein-AM level than the *ΔcyoB E. coli*-treated N2 wild-type worms and the BW25113-treated *bas-1(lf)* worms (Fig. 3A and 3B), indicating the further reduction of labile iron caused by the *bas-1(lf)* in the presence of *ΔcyoB E. coli*. These results together indicate the function of elevating levels of dopamine and serotonin in the neurons for attenuating bacteria-induced reductions in labile iron levels.

**Fig. 3:**
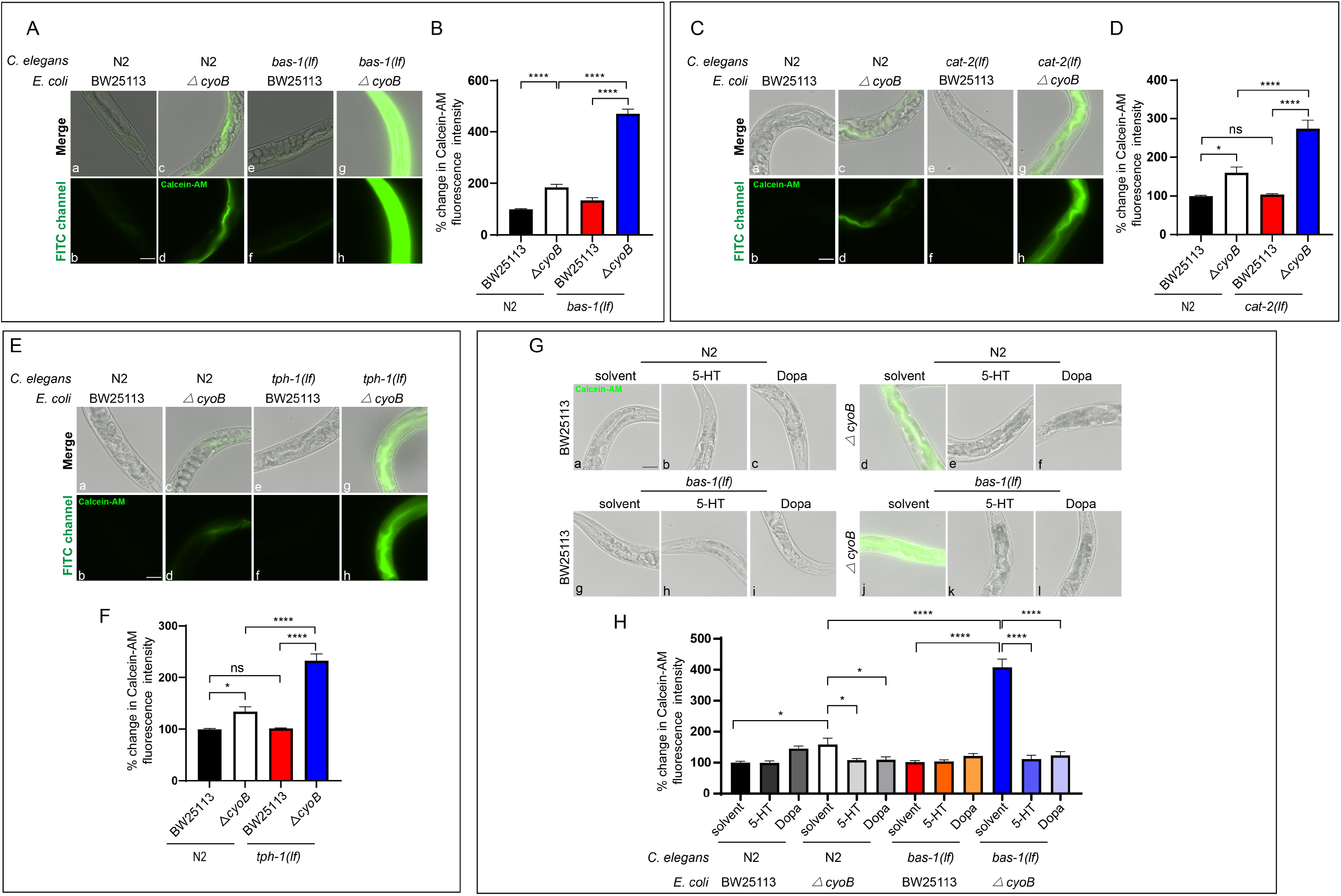
The increase in *bas-1* expression in serotonergic and dopaminergic neurons in response to the presence of *ΔcyoB E. coli* is crucial for the *C. elegans* host to alleviate the resultant reduction in labile iron. **(A and B)** Images and bar graph shows the effects of inhibiting dopamine and serotonin biosynthesis enzyme *bas-1* on labile iron level in the presence of BW25113 and *ΔcyoB E. coli*. The percent change in Calcein fluorescence was calculated by normalization to the levels in N2 worms treated with BW25113. **(C and D)** Images and bar graph showing that deleting dopamine biosynthesis enzyme *cat-2* aggravates the reduction in labile iron caused by *ΔcyoB E. coli*. Calcein-AM was used to indicate the labile iron and the % change in Calcein fluorescence was calculated by normalization to the levels in N2 worms treated with BW25113. **(E and F)** Images and bar graph illustrating the effect of *tph-1* deletion on the reduction in labile iron levels at the presence of the indicated bacteria. TPH-1 is essential for serotonin biosynthesis, as shown in Figure 1A. The percent of change in Calcein fluorescence intensity relative to the levels in N2 worms treated with BW25113 was quantified. **(G and H)** Images and bar graph indicating that the supplementation of 12.5 mg/ml dopamine (Dopa) and 10 mg/ml serotonin (5-HT) is able to reverse the *ΔcyoB E. coli*-caused free iron reduction in the wild-type N2 worms as well as the exacerbated labile iron reduction in the *bas-1(lf)* animals treated with *ΔcyoB E. coli*. The % change in the Calcein fluorescence intensity relative to the levels in the wild type N2 worms treated with both BW25113 and solvent is shown in the bar graph. For all the panels, experiments were performed for at least three times and data in the bar graph are the mean ± SEM. *p < 0.05 and ****p < 0.0001 by one-way ANOVA; Scale bar, 50 μm. n > 25 each group.

To determine which of these two types of neurons mediated this host response, we tested whether *C. elegans* strains with single deletions of *tph-1* (*tph-1(lf)*) and *cat-2* (*cat-2(lf)*), which show disrupted serotonin and dopamine biosynthesis, respectively ^25^, display exacerbated reductions in labile iron levels following treatment with *ΔcyoB E. coli*. Interestingly, Calcein-AM analysis indicated that both *cat-2(lf)* and *tph-1(lf)* animals displayed an exacerbated reduction in the levels of labile iron in the presence of *ΔcyoB E. coli* (Fig. 3C g-h and 3D; Fig. 3E g-h and 3F), in contrast to the *ΔcyoB E. coli*-treated N2 wild-type worms, which displayed a less severe reduction in labile iron (Fig. 3C c-d and 3D; Fig. 3E c-d and 3F), and *cat-2(lf)* and *tph-1(lf)* mutant worms treated with BW25113, which showed no obvious reduction in labile iron (Fig. 3C e-f and 3D; Fig. 3E e-f and 3F). However, in the presence of *ΔcyoB E. coli*, *tph-1(lf)* and *cat-2(lf)* mutations did not cause as severe a reduction in labile iron levels as that caused by the *bas-1(lf)* mutation (Fig. 3 A-F), which inhibits both dopamine and serotonin biogenesis, suggesting that serotonergic and dopaminergic neurons function redundantly to promote labile iron levels. Therefore, when confronting bacteria-caused reductions in labile iron levels, the responses of both the serotonergic and dopaminergic neurons are required for the host to maintain iron homeostasis.

Next, we sought to determine whether serotonergic and dopaminergic neurons modulate iron homeostasis through serotonin and dopamine signaling; thus, we performed dopamine and serotonin supplementation to analyze their potential effects on alleviating *bas-1(lf)*-caused aggravated labile iron reduction. We observed that supplementation with either serotonin or dopamine was able to almost fully reverse the reduction in iron levels in the *bas-1(lf)* mutants treated with *ΔcyoB E. coli* (Fig. 3G g-l and 3H), which indicates that *bas-1*-expressing neurons function through serotonin and dopamine neurotransmitters to regulate iron metabolism. Given that increasing the bioavailability of neurotransmitters by supplementation can over-activate the corresponding signaling pathways ^31^, these results support that over-activating either dopamine or serotonin signaling could overcome the iron reduction caused by the inhibition of both serotonin and dopamine signaling in the *bas-1(lf)* worms, respectively. Corroborating this idea, we observed that either dopamine or serotonin supplementation could even suppress *ΔcyoB E. coli*-induced reductions in labile iron levels in the wild-type worms (Fig. 3G a-f and 3H), thus overactivation of either dopamine or serotonin signaling is capable of elevating labile iron levels.

To further confirm that dopaminergic and serotonergic neurons regulate iron metabolism through dopamine and serotonin signaling, we also analyzed a potential role of *cat-1*, encoding transporter for loading serotonin and dopamine into the vesicle for release (Fig. 1A) ^25^, in facilitating worms to cope with bacteria-induced reduction of available iron. Indeed, we found that, in the presence of *ΔcyoB E. coli,* inhibiting the release of both dopamine and serotonin by deleting *cat-1(lf)* caused an aggravated reduction in labile iron levels (Supplementary information, Fig. S3 A-B), when compared to the labile iron levels in the BW25113 treated*-cat-1(lf)* worms and the *ΔcyoB E. coli-*treated N2 wild-type worms. The level of labile iron in the *ΔcyoB E. coli* treated*-cat-1(lf)* worms is similar to that in the *ΔcyoB E. coli* treated*-bas-1(lf)* worms (Fig. 3 A-B and Supplementary information, Fig. S3 A-B), consistent with that CAT-1 and BAS-1 are required for both dopamine and serotonin signaling. In summary, dopaminergic and serotonergic neurons inspect bacteria-induced reductions in labile iron levels and then orchestrate the host response through dopamine and serotonin signaling to counteract this decrease.

### Head serotonergic and dopaminergic neurons respond to the bacteria*-*induced labile iron reduction by perceiving the consequent impairment of the intestinal mitochondria, and this neuronal response is crucial for the host to maintain mitochondrial function when confronting these bacteria

We next aimed to dissect how dopaminergic and serotonergic neurons perceive changes in labile iron levels by determining whether this process involved mitochondria, given that cellular respiration in mitochondria is one of the major processes that utilizes iron ^32^. To this end, we tested the possibility that these neurons perceive the reduction in labile iron levels by monitoring mitochondrial impairment.

First, we measured the oxygen consumption rate (OCR) to assess the change in mitochondrial function after bacterial insult. Our analyses showed that the OCR was lower in the wild-type worms treated with *ΔcyoB E. coli* than in the ones treated with BW25113 by almost 2-fold (Fig. 4A). Interestingly, this reduction in OCR caused by *ΔcyoB E. coli* could be reversed by iron supplementation (Fig. 4A). Therefore, *ΔcyoB E. coli*-induced labile iron insufficiency causes a decline in mitochondrial function. To probe a possible role of the mitochondrial decline in triggering response in the dopaminergic and serotonergic neurons, we next induced mitochondrial impairment by inhibiting the function of SPG-7 mitochondrial metalloprotease that has been widely used to induce mitochondrial stress ^33^ and evaluated a potential induction of the BAS-1 expression in neurons. Strikingly, the increase in BAS-1::GFP expression was induced by *spg-7(RNAi)* (Fig. 4B and 4C), which indicates a causal role of the decline in mitochondrial function in triggering the response of dopaminergic and serotonergic neurons to the *△cyoB E. coli-*induced decrease in iron levels. Therefore, the dopaminergic and serotonergic neurons respond to the *△cyoB E. coli-*caused iron decrease by perceiving the consequent mitochondrial impairment.

**Fig. 4:**
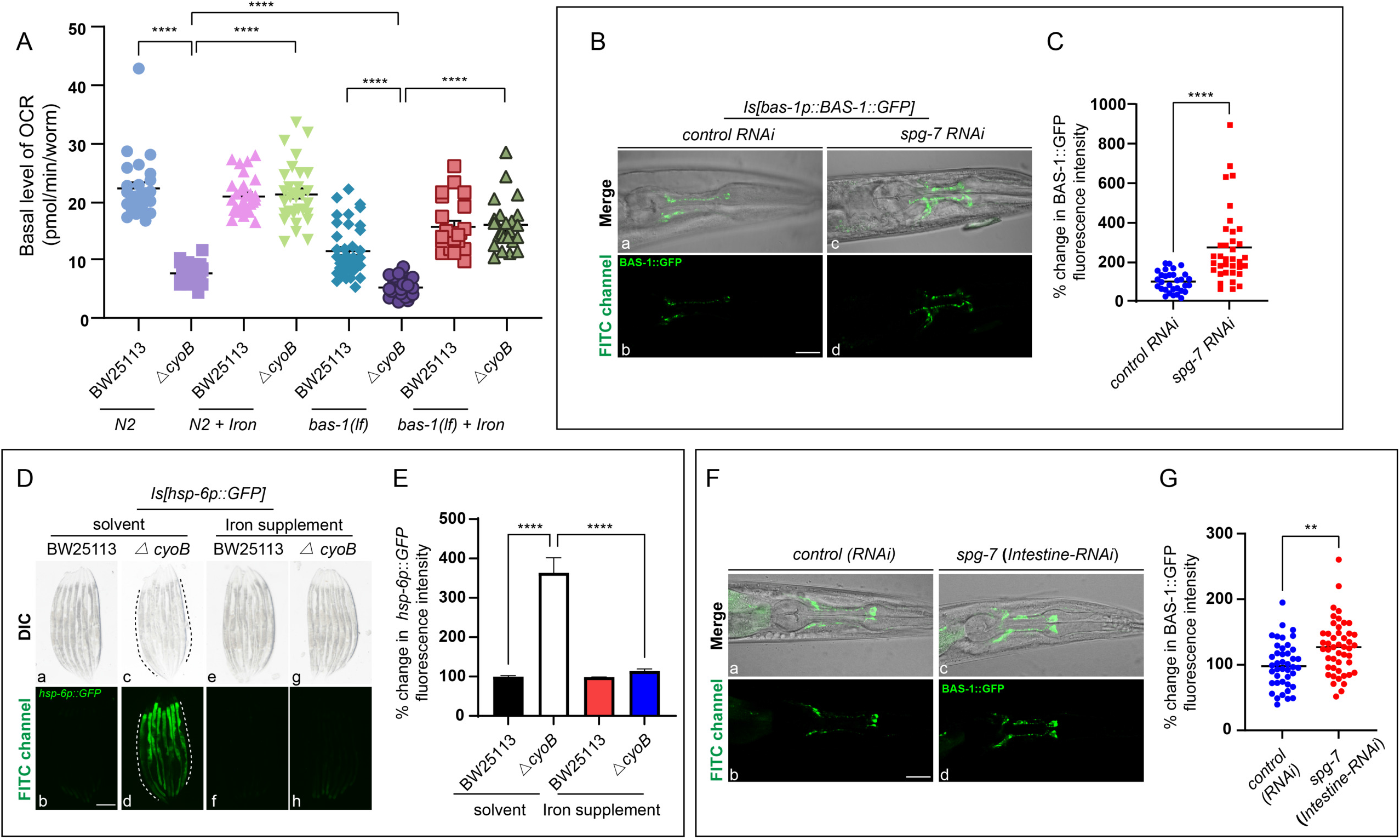
Serotonergic and dopaminergic neurons respond to Δ*cyoB E. coli*-induced reductions in labile iron levels by perceiving the impairment of intestinal mitochondria, and this neuronal response is crucial for coping with the bacteria-caused reduction in mitochondrial function. **(A)** Graphs showing the reduction in the basal oxygen consumption rate (OCR) in wild-type animals treated with *ΔcyoB E. coli* and the further reduction in the OCR in dopamine- and serotonin-deficient animals incubated with *ΔcyoB E. coli*. Iron supplementation was performed by adding 4mM FeCl_3_ to the worm culture medium. The OCR was analyzed by a Seahorse XFe96 Analyzer as previously shown ^45^, and each point indicates the OCR normalized to each worm. ****p < 0.0001 by unpaired t-test; n > 100 for each treatment. **(B and C)** Representative images and plotted graph indicating that the decline in mitochondrial function caused by *spg-7(RNAi)* is able to trigger the expression of BAS-1::GFP in head neurons. Each dot represents the % change in BAS-1::GFP calculated by normalization to the levels in worms treated with control RNAi. ****p < 0.0001 by unpaired t test. n > 30 each group. Scale bar, 20 μm. **(D and E)** Microscopic images and bar graph showing the effects of the *ΔcyoB* bacteria on mitochondrial damage with or without iron supplementation. The induction of the expression of *hsp-6p::GFP*, a widely used mitochondrial stress reporter, was employed to indicate mitochondrial damage. Dashed lines indicate the intestine and 4mM FeCl_3_ was added to the worms carrying *hsp-6p::GFP* for iron supplementation. The % change in *hsp-6p::GFP* was calculated by normalization to the levels in worms treated with BW25113. Data are the mean ± SEM. ****p < 0.0001 by one-way ANOVA; n > 20 each group. Scale bar, 200 μm. **(F and G)** Microscopic images and graph indicating that *spg-7(intestinal RNAi)* is able to elevate BAS-1::GFP expression. The *rde-1(lf); kbIs7 [nhx-2p::RDE-1+rol-6(su1006)]; Is[bas-1p::BAS-1::GFP]* worm strain was used to perform *spg-7(intestine-RNAi)* for inducing intestinal mitochondrial impairment, and then analyze the resultant impact on the BAS-1 expression. The animals treated with the HT115 *E. coli* strain carrying the empty PL4440 vector served as the RNAi control to calculate the percent change of BAS-1::GFP fluorescence intensity. Each dot represents the percentage change of BAS-1::GFP intensity of each worm relative to the average BAS-1::GFP intensity in *control RNAi*-treated worms. **p < 0.01 by unpaired t test. n > 30 each group. Scale bar, 20 μm.

We then aimed to determine whether the dopaminergic and serotonergic neurons perceive the decline in mitochondrial function in peripheral tissue or in neurons. First, we analyzed in which *C. elegans* tissue the mitochondria damage was caused by the *ΔcyoB E. coli* by evaluating the expression of the commonly-used mitochondrial stress reporter Is[*hsp-6p::GFP*], which can be induced in neurons as well as peripheral tissues when mitochondria are impaired in corresponding cells ^33^.

Intriguingly, the obvious induction of *hsp-6p::GFP* expression in the intestinal cells, but not in the neurons, was observed in the *ΔcyoB E. coli-*treated worms (Fig. 4 D-E and Fig. S4A). Moreover, this induction of *hsp-6p::GFP* in *ΔcyoB E. coli-*treated worms could be suppressed by iron supplementation (Fig. 4 D-E). These results indicate that *ΔcyoB E. coli-*induced decreases in labile iron levels resulted in mitochondrial impairment mainly in the intestine. Next, we tested the possibility that head dopaminergic and serotonergic neurons may sense the reduction in iron levels by perceiving this decline in mitochondrial function in the gut. To this end, we determined whether inducing mitochondrial impairment specifically in the gut with intestine-specific RNAi against *spg-7* could increase BAS-1 expression in neurons. We performed *spg-7 RNAi* in the *Is[bas-1p::BAS-1::GFP]*; *rde-1(lf); kbIs7 [nhx-2p::RDE-1+rol-6(su1006)]* worms, in which the RNAi is only effective in the intestine ^34^. Indeed, we found that *spg-7 (*intestine*-RNAi)* led to significantly higher BAS-1 expression than the control RNAi (Fig. 4 F-G), indicating that BAS-1-expressing neurons monitor intestinal mitochondrial function state. Moreover, reducing labile iron levels by BP treatment caused a decline in mitochondrial function in the gut, as evidenced by the expression of *hsp-6p::GFP* in the intestine (Supplementary information, Fig. S4B); additionally, expression of BAS-1::GFP was induced by BP as described above (Fig. 2 E-F), which further supports the idea that reduced labile iron levels caused a decline in intestinal mitochondrial function and triggered BAS-1 expression in dopaminergic and serotonergic neurons. In summary, host neurons perceive *ΔcyoB E. coli*-induced labile iron insufficiency by detecting intestinal mitochondrial impairment to trigger neuronal BAS-1 expression to orchestrate a response to recover the labile iron level.

Based on these data, we hypothesized that the response that enabled these neurons to attenuate the reduction in labile iron levels might be important for mitigating *ΔcyoB E. coli-*caused impairment of mitochondrial function. Consistent with our prediction, *bas-1(lf)* animals treated with *ΔcyoB E. coli* displayed even lower OCR than the *bas-1(lf)* animals treated with BW25113 and the wild-type worms cultured with *ΔcyoB E. coli* (Fig. 4A), indicating that the neuronal response is important for maintaining mitochondrial function when the *ΔcyoB E. coli* are present. Furthermore, the further reduction in OCR in the *bas-1(lf)* worms treated with *ΔcyoB E. coli* was obviously suppressed by iron supplementation (Fig. 4A). Together with the observation that the *bas-1(lf)* mutation aggravated the *ΔcyoB E. coli*-induced reduction in labile iron levels, these results indicate that in response to *△cyoB E. coli*, the upregulation of BAS-1 expression to increase dopamine and serotonin levels in these neurons plays a role in improving mitochondrial function at least partially by promoting available iron level.

To further investigate the important role of the induction of *bas-1* expression in neurons in maintaining mitochondrial function under conditions of iron scarcity, we tested the possibility that inhibiting *bas-1* expression would impair the ability of *ΔcyoB E. coli*-treated worms to deal with mitochondrial stress. Paraquat, a widely used mitochondrial stressor ^35^, was employed to induce mitochondrial dysfunction.

After paraquat treatment, the survival time of wild-type worms cultured with *ΔcyoB E. coli* was lower than that of worms treated with BW25113, which is in agreement with the decrease in the labile iron level and decline of mitochondrial function in these wild-type worms treated with *ΔcyoB E. coli* (Supplementary information, Fig. S4C). Importantly, this survival rate reduction could be fully reversed by iron supplementation (Supplementary information, Fig. S4C), which is in agreement with the mitochondrial function decline caused by the *ΔcyoB E. coli-*induced reduction in labile iron levels. These results indicate that *ΔcyoB E. coli*-induced reductions in labile iron levels impairs the host’s ability to cope with mitochondrial stress.

Furthermore, consistent with our prediction, paraquat treatment induced an even lower survival rate in the *bas-1(lf)* mutants treated with *ΔcyoB E. coli* than in the wild-type worms treated with *ΔcyoB E. coli* or the *bas-1(lf)* mutants cultured with BW25113 wild-type *E. coli*. This *bas-1(lf)*-induced further decrease in the survival rate could be strongly reversed by iron supplementation (Supplementary information, Fig. S4C). Altogether, these data suggest that, in the presence of bacteria that induce reductions in available iron levels, triggering a response from dopaminergic and serotonergic neurons to recover labile iron levels is crucial for improving animals’ ability to maintain mitochondrial homeostasis.

### The mitochondrial stress regulator ATFS-1 acts in the intestine to mediate dopaminergic and serotonergic neuron perception of *ΔcyoB E. coli-*induced reductions in labile iron levels

In order to understand how neurons perceive the intestinal mitochondrial impairment caused by bacteria-induced iron insufficiency, we next aimed to dissect the molecular mechanisms underlying this gut-brain communication. As ATFS-1, expressed in the majority of cells and tissues including the intestine in *C. elegans* ^36^, is known to be a master regulator for sensing mitochondrial dysfunction ^37^, we thus evaluated whether intestinal ATFS-1 might be required for transmitting intestinal mitochondrial damage signals to *bas-1*-expressing neurons. We performed *atfs-1 RNAi* in the *Is[bas-1p::BAS-1::GFP]*; *rde-1(lf); kbIs7 [nhx-2p::RDE-1+rol-6(su1006)]* worms for specifically knocking down *atfs-1* in the intestine ^34^. Indeed, we found that, the higher BAS-1::GFP in the worms treated with the *ΔcyoB E. coli* than the ones treated with the wildtype bacteria treatment (Fig. 5A, 5B a-d and 5C), was suppressed by the intestine-specific knockdown of *atfs-1* when compared to the treatment with the RNAi control (Fig. 5B c-d/g-h and 5C), which indicates that intestinal ATFS-1 is required for the response of dopaminergic and serotonergic neurons to the mitochondrial impairment in the gut. Furthermore, we tested whether activating ATFS-1 in the intestine via a transgene expressing the dominant-active form of ATFS-1 with intestine-specific promoter *gly-19* ^33,37^ was able to induce *Is[bas-1p::BAS-1::GFP]* in the presence of wildtype bacteria. Strikingly, compared to the no transgene control treatment, expression of this *atfs-1(intestine-gf)* transgene was able to strongly increase neuronal BAS-1 expression (Fig. 5D and 5E). Altogether, these data indicate that ATFS-1 acts in the intestine to perceive intestinal mitochondrial impairment and then triggers BAS-1 expression in neurons, thus mediating the impact of *ΔcyoB E. coli* on dopaminergic and serotonergic neurons.

**Fig. 5:**
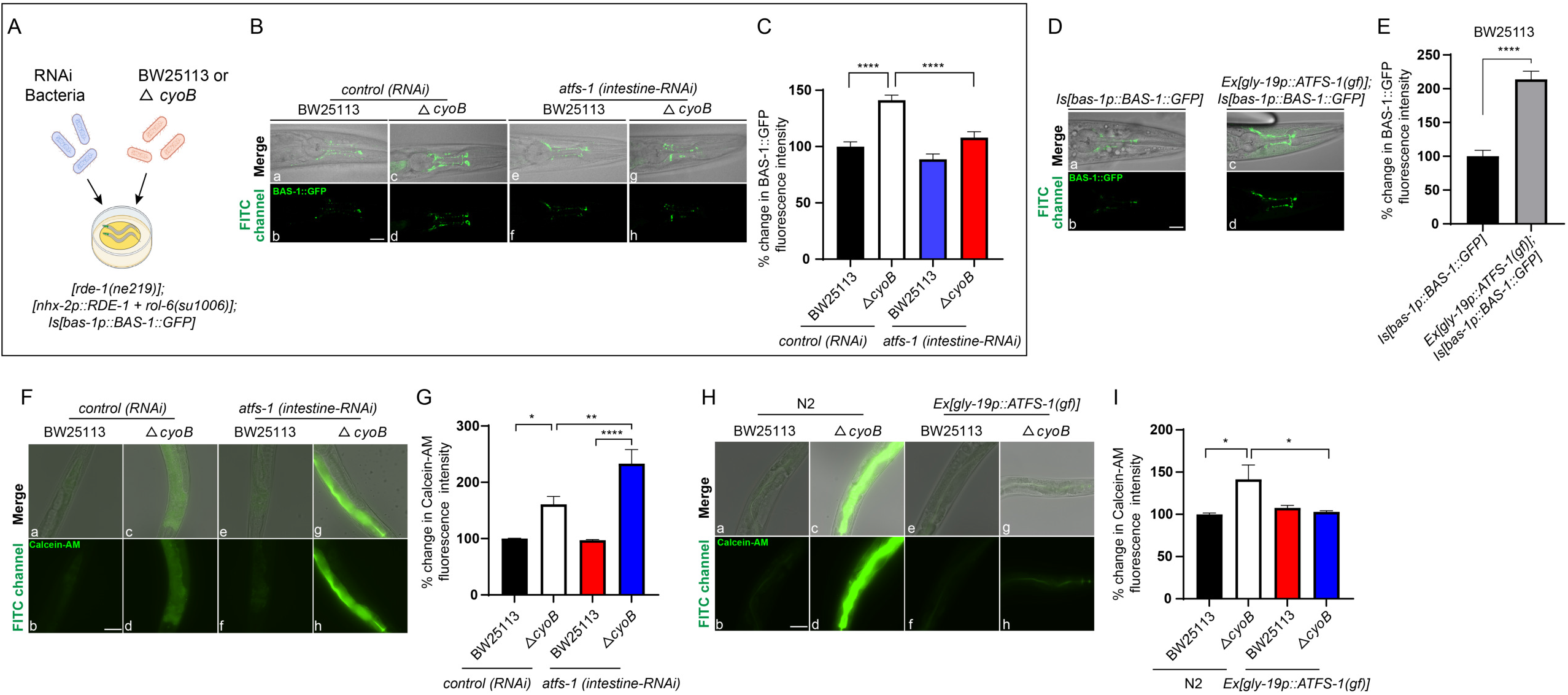
The intestine-expressed mitochondrial stress sensor ATFS-1 mediates the response of dopaminergic/serotonergic neurons to Δ*cyoB E. coli* to attenuate reductions in labile iron levels. **(A-C)** Images and bar graphs showing that intestinal ATFS-1 is required for the *ΔcyoB E. coli*-induced increase in neuronal *bas-1* expression. BW25113 or *ΔcyoB E. coli* was mixed with *atfs-1* RNAi-containing bacteria to analyze the role of *atfs-1* in mediating the bacteria-triggered response as described in (A). The *rde-1(lf); kbIs7 [nhx-2p::RDE-1+rol-6(su1006)]* worm strain expressing BAS-1::GFP underwent knock down of *atfs-1* specifically in the intestine. The % change in BAS-1::GFP fluorescence was calculated by normalization to the levels in animals treated with control RNAi and BW25113. Statistics analyzed by one-way ANOVA; Scale bar, 20 μm. n > 50 each group. **(D and E)** Images and bar graph showing that overexpression of the dominant active form of ATFS-1 in the intestine induces neuronal expression of BAS-1::GFP. The intestine-specific promoter *gly-19* was used to drive the intestinal expression of *atfs-1(gf)* ^33^. The worms with indicated genotypes were treated with BW25113. The % change in BAS-1::GFP fluorescence intensity was calculated by normalization to the levels in the worms without transgene. Statistics analyzed by unpaired t test. Scale bar, 20 μm. n > 30 each group. **(F and G)** Images and bar graph indicating that inhibiting intestinal *atfs-1* expression exacerbates the reduction in labile iron caused by *ΔcyoB* bacteria. The *rde-1(lf); kbIs7 [nhx-2p::RDE-1+rol-6(su1006)]* worm was used for *atfs-1* intestine-specific RNAi analyses. As performed in (A), the RNAi bacteria were mixed with the BW25113 or *ΔcyoB E. coli* to analyze the role of intestinal *atfs-1* in coping with bacteria-caused reductions in iron levels. Calcein-AM analyses were used to indicate the level of labile iron. The % change in Calcein fluorescence was calculated by normalization to the levels in animals treated with control RNAi and BW25113. Statistics analyzed by one-way ANOVA; n > 20 each group. Scale bar, 50 μm. **(H and I)** Images and bar graphs showing that intestine-specific overexpression of dominant active form of ATFS-1 is able to recover the *ΔcyoB* bacteria-caused labile iron reduction in the N2 wildtype worms. The *gly-19* promoter was used to drive the expression of *atfs-1(gf)* specifically in the intestine. The % change of Calcein fluorescence was determined by normalization to the BW25113-treated N2 wild-type worms. Statistics analyzed by one-way ANOVA; n > 20 each group. Scale bar, 50 μm. Fall the panels, experiments were performed for at least three times. Data are the mean ± SEM. *p < 0.05, **p < 0.01 and ****p < 0.0001.

Furthermore, we aimed to investigate whether intestinal ATFS-1-mediated induction of neuronal BAS-1 is involved in orchestrating response to modulate iron metabolism as the neuronal responses triggered by the *ΔcyoB E. coli* bacteria. First, we analyzed whether knock-down of *atfs-1* specifically in the intestine aggravated iron reduction in the presence of *ΔcyoB E. coli*. Indeed, *atfs-1(intestinal RNAi)* worms treated with *ΔcyoB E. coli* displayed even greater reduction in labile iron levels than the *atfs-1(intestinal RNAi)* worms treated with wildtype *E. coli* or the animals treated with both control RNAi and *ΔcyoB E. coli* (Fig. 5F and 5G), which was similar to the findings observed in the *bas-1(lf)* group (Fig. 3A and 3B). Conversely, intestine-specific expression of the dominant-active form of ATFS-1 was able to fully suppress the *ΔcyoB E. coli-*induced reduction in labile iron levels in the wildtype N2 worms (Fig. 5H and 5I), which was also observed in the *atfs-1(gf)* mutant worms (Supplementary information, Fig. S5A and S5B), indicating that ATFS-1 mainly functions in the intestine to cope with *ΔcyoB E. coli*-caused labile iron reduction. In summary, intestinal *atfs-1* senses the iron insufficiency-induced impairment of intestinal mitochondria to promote BAS-1 expression in dopaminergic and serotonergic neurons to in turn alleviate intestinal iron deficiency, thus mediating the perception of *ΔcyoB E. coli* by the dopaminergic and serotonergic neurons.

### Neuronal CWN-2 Wnt mediates the role of intestinal ATFS-1 in elevating *bas-1* expression in dopaminergic and serotonergic neurons to increase labile iron levels

Our data suggested that cell non-autonomous signaling may mediate communication between the intestinal cells and *bas-1*-expressing neurons to modulate iron metabolism. Discovering the cell non-autonomous signal that regulates iron metabolism is highly important, given that maintaining systemic iron homeostasis is critical for organism health. Thus, we next explored whether any endocrine signals might be involved in regulating *ΔcyoB E. coli*-induced changes in iron metabolism, and genes, encoding ligands and receptors that are involved in several major signaling pathways, were tested by analyzing the impact of their deletions on labile iron level in the presence of *ΔcyoB E. coli* (Supplementary information, Fig. S6A). Interestingly, *cwn-2(lf)* worms treated with *ΔcyoB E. coli* displayed even lower reduction in labile iron than in the *cwn-2(lf)* worms treated with *wildtype E. coli or in the* N2 wildtype worms treated with *ΔcyoB E. coli* (Fig. 6A). Therefore, deletion of the CWN-2 Wnt ligand exacerbated the reduction in labile iron levels caused by *ΔcyoB E. coli*, which was similar to the effect observed in the *bas-1(lf)* mutant worms (Fig. 3 A-B), suggesting a potential increase of CWN-2 in response to *ΔcyoB E. coli* for alleviating the iron insufficiency. Thus, we next analyzed the transcription of *cwn-2* by qPCR in the N2 worms treated with *ΔcyoB E. coli* and BW25113 wild type bacteria. As predicted, we found that the transcription level of *cwn-2* was increased by *ΔcyoB E. coli* treatment (Fig. 6B), and this increase was entirely suppressed by iron supplementation (Fig. 6B). These results together indicate that *cwn-2* levels are increased in response to labile iron insufficiency caused by *ΔcyoB E. coli* to counteract further reductions in iron levels.

**Fig. 6:**
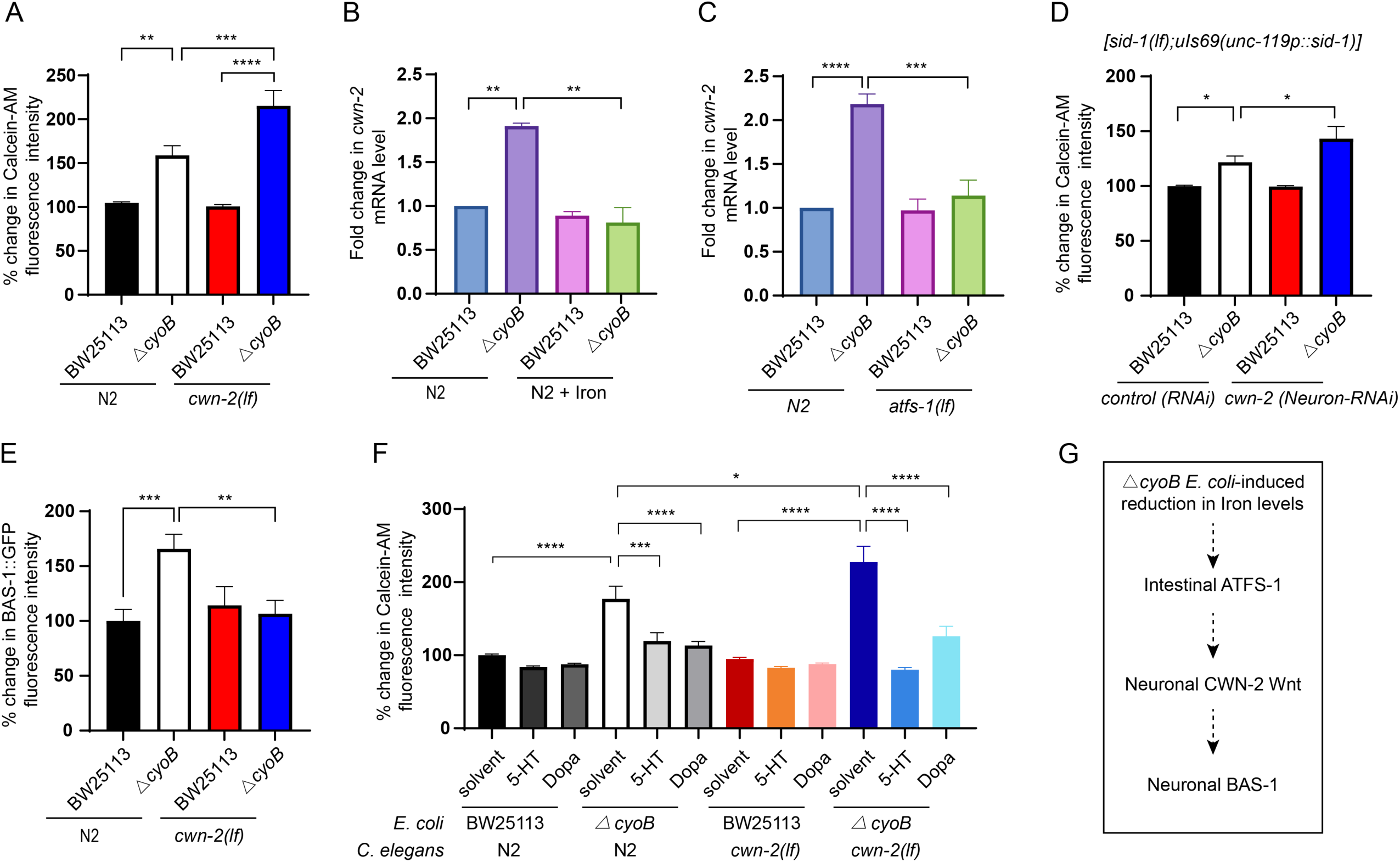
Neuron-expressed CWN-2 Wnt acts downstream of ATFS-1 to mediate the response of dopaminergic/serotonergic neurons to *ΔcyoB E. coli*-caused labile iron insufficiency. **(A)** Bar graph showing the effect of CWN-2 Wnt ligand mutation on *ΔcyoB E. coli*-caused reduction in labile iron levels analyzed by Calcein-AM staining. The data are normalized to Calcein fluorescence in wild-type N2 *C. elegans* treated with BW25113 bacteria. n > 50 each group. **(B)** qPCR measurements of *cwn-2* mRNA levels indicating that *ΔcyoB E. coli* bacteria-caused labile iron insufficiency induced the transcription of *cwn-2* in *C. elegans*. 4mM FeCl_3_ was used for iron supplementation. Fold change in *cwn-2* mRNA levels was analyzed by normalizing to the levels in BW25113-treated animals. n > 200 each group. **(C)** qPCR measurements of *cwn-2* mRNA levels showing that the increase in *cwn-2* transcription caused by *ΔcyoB E. coli* is dependent on *atfs-1*. The fold change in the levels in animals subjected to each treatment was calculated by normalizing to the levels in wild-type animals treated with BW25113 bacteria. n > 200 each group. **(D)** Bar graph showing that knocking down *cwn-2* in neurons exacerbates the *ΔcyoB E. coli*-induced reduction in labile iron in the *C. elegans* host. The neuron-specific RNAi strain TU3401[*sid-1(lf);uIs69(unc-119p::sid-1)*] was employed to knock down neuronal *cwn-2.* n > 20 for each group. **(E)** Bar graph indicating that *cwn-2* is required for the *ΔcyoB E. coli-*triggered increase in BAS-1 expression. *Is[bas-1p::BAS-1::GFP]* and *cwn-2(lf);Is[bas-1p::BAS-1::GFP]* animals were treated with the indicated bacteria. The fold change in BAS-1::GFP fluorescence intensity was calculated by normalization to the *Is[bas-1p::BAS-1::GFP]* animals treated with BW25113. n > 30 for each treatment. **(F)** Bar graph indicating that *cwn-2* functions upstream of dopamine and serotonin signaling to attenuate bacteria-caused iron reduction analyzed with Calcein-AM staining. Supplementation with 12.5 mg/ml dopamine (Dopa) or 10 mg/ml serotonin (5-HT) was able to recover the labile iron level in N2 and *cwn-2(lf)* worms treated with *ΔcyoB E. coli*. An equivalent volume of water was used as the solvent control. n > 30 for each treatment. **(G)** A schematic depicting the functional relationship of the indicated proteins in the process through which dopaminergic and serotonergic neurons perceive the presence of *ΔcyoB E. coli*. For all the panels, experiments were performed for at least three times and data are the mean ± SEM. *p < 0.05, **p < 0.01, ***p < 0.001 and ****p < 0.0001 by one-way ANOVA.

Next, we investigated the functional relationship between *cwn-2* and *atfs-1.* We found that the *ΔcyoB E. coli-*induced increase in *cwn-2* transcription was suppressed by the *atfs-1(lf)* mutation (Fig. 6C), indicating that CWN-2 acts downstream of ATFS-1 to respond to *ΔcyoB E. coli*-induced labile iron insufficiency. Furthermore, we performed tissue-specific RNAi analyses to investigate whether the CWN-2 Wnt ligand originates from the intestine or from neurons to act downstream of ATFS-1 to modulate iron metabolism. Our results indicated that knocking down *cwn-2* specifically in the intestine did not exacerbate the reduction in labile iron levels in the animals treated with *ΔcyoB E. coli* (Supplementary information, Fig. S6B). In contrast, when performing the neuron-specific RNAi against *cwn-2* by using the *sid-1(lf);uIs69(unc-119p::sid-1)* strain, a further reduction in labile iron levels occurred in the presence of *ΔcyoB E. coli* (Fig. 6D). The less severe a reduction in the labile iron in the *cwn-2(Neuron-RNAi)* worms treated with *ΔcyoB E. coli* than in the *cwn-2(lf)* worms treated with *ΔcyoB E. coli* may be due to the partial knock-down of *cwn-2* by RNAi. These results support that the neuron-originated CWN-2 Wnt ligand, that responds to the intestinal ATFS-1, is required for the host to orchestrate a response to *ΔcyoB E. coli* challenge to attenuate reductions in available iron levels.

In addition, in our candidate screen shown in Figure S6A, we found that *egl-20*, another Wnt ligand, was also required for alleviating *ΔcyoB E. coli-*induced iron reduction (Supplementary information, Fig. S6C), confirming the unexpected role of Wnt signaling in modulating iron metabolism. In response to the presence of *ΔcyoB E. coli,* the increased transcription of *egl-20* was less obvious than that of *cwn-2* (Supplementary information, Fig. S6D; Fig. 6B); thus, we focused on the role of *cwn-2* in triggering the neuronal response in this study. Interestingly, from our screen (Supplementary information, Fig. S6A), we found that inhibiting the insulin pathway was able to recover the labile iron level in worms treated with *ΔcyoB E. coli* (Supplementary information, Fig. S6A), suggesting a repressing role of insulin-signaling in regulating labile iron level. It is worth testing the role of the insulin pathway in mediating bacteria-host iron talk in the future. Altogether, our data support that Wnt signaling originating in neurons acts downstream of intestinal ATFS-1 to mediate the host response to *ΔcyoB E. coli* for counteracting bacteria-induced reductions in labile iron levels.

Our data above indicated that both *bas-1* and *cwn-2* were expressed in neurons to positively regulate labile iron levels. Thus, we next evaluated the functional relationship between CWN-2 and serotonin and dopamine signaling. We found that the *ΔcyoB E. coli*-induced expression of BAS-1 was repressed by the deletion of *cwn-2* (Fig. 6E). In contrast, the *bas-1(lf)* mutation showed no obvious effect on *cwn-2* expression induced by *ΔcyoB E. coli* (Supplementary information, Fig. S6E). These results indicate that *cwn-2* acts upstream of *bas-1* and downstream of intestinal ATFS-1 to mediate the dopaminergic and serotonergic neuron response to the presence of *ΔcyoB E. coli*. Furthermore, overactivation of dopaminergic and serotonergic signaling by supplementing dopamine and serotonin, respectively, not only strongly suppressed the reduction in iron levels in the wild-type worms cultured with *ΔcyoB E. coli* but also almost entirely reversed the *cwn-2(lf)*-induced further reduction in labile iron levels in the presence of *ΔcyoB E. coli* (Fig. 6F). This result further demonstrates that BAS-1-involved dopamine and serotonin biosynthesis act to mediate the role of CWN-2 Wnt in promoting labile iron level. Overall, in the presence of *ΔcyoB E. coli, bas-1* expression in dopaminergic and serotonergic neurons is triggered by a *cwn-2* Wnt signal originating in neurons to elevate peripheral labile iron levels for alleviating the bacteria-induced reductions in iron levels (Fig. 6G).

### The increase in BAS-1 expression in head dopaminergic and serotonergic neurons attenuates bacteria*-*induced reduction in labile iron levels by promoting intestinal *ferritin-1(ftn-1)* expression, which indicates an unexpected gut-brain-gut axis that enables the host to cope with bacteria-caused reductions in available iron

Next, we aimed to investigate the molecular mechanism by which the induction of *bas-1* expression in dopaminergic and serotonergic neurons attenuates the reduction in labile iron levels. Since cellular labile iron levels can be elevated by increasing iron absorption, we first tested whether the gene expression of the iron transporters SMF-1/-2/-3 for absorbing iron is mediated by neuronal BAS-1 after bacteria treatment. Our results indicated that compared to the wild type bacteria control, the *ΔcyoB E. coli* treatment increased the transcription of *smf-1/-2/-3* (Supplementary information, Fig. S2 and S7 A-C); however, we found that deletion of *bas-1* did not affect the induction of *smf-1/-2/-3* expression by *ΔcyoB E. coli* treatment (Supplementary information, Fig. S7 A-C), indicating that the *ΔcyoB E. coli*-induced expression of *smf-1/-2/-3* is not dependent on the activation of *bas-1*-expressing neurons. Therefore, in the presence of *ΔcyoB E. coli,* the dopaminergic and serotonergic neurons are unlikely to elevate iron levels by increasing the expression of these iron transporters.

Iron storage is known to repress labile iron levels through iron sequestration by ferritin, and ferritin levels have been previously shown to be reduced in response to iron deficiency to release labile iron ^29^. We thus tested whether dopaminergic and serotonergic neurons might elevate the free iron level by decreasing iron storage protein ferritin. We performed qPCR analyses of the mRNA level of *ftn-1*, a critical iron storage factor whose expression was reported to be transcriptionally repressed at iron-scarce conditions ^29^, in the N2 wildtype animals treated with BW25113 and *ΔcyoB E. coli.* Surprisingly, we found that compared to the BW25113 parent bacteria control treatment, *ΔcyoB E. coli* that displayed strong iron reducing effect obviously increased rather than decreased *ftn-1* transcription (Fig. 7A). Furthermore, analysis of a *ftn-1* transcriptional reporter (*ftn-1p::*GFP) confirmed this obvious increase in *ftn-1* transcription in response to the presence of *ΔcyoB E. coli* (Fig. 7 B-C). Specifically, we observed that *ftn-1* displayed intestine-specific expression as previously described (Fig. 7B a-b and Supplementary information, Fig. S7D) ^29,38^, and after *ΔcyoB* treatment, a clear increase in *ftn-1p::GFP* expression in the intestine was observed (Fig. 7B c-d and Supplementary information, Fig. S7D). Therefore, these results suggest that this bacteria-induced decrease in labile iron level boosts *ftn-1* expression. Further confirming this idea, *ΔcyoB E. coli-*induced *ftn-1* mRNA (Fig. 7A) and *ftn-1p::GFP* expression in the intestine (Fig. 7B e-h and 7C) were both abolished by iron supplementation. These results together demonstrate that *ftn-1* transcription was increased in response to *ΔcyoB E. coli-*induced labile iron insufficiency. Importantly, we observed that the induction of intestinal *ftn-1* expression by *ΔcyoB E. coli* was partially inhibited by the *bas-1(lf)* mutation (Fig. 7D). Therefore, our findings indicate that dopaminergic and serotonergic neurons promote intestinal *ftn-1* transcription in response to *ΔcyoB E. coli*-induced labile iron reduction.

**Fig. 7:**
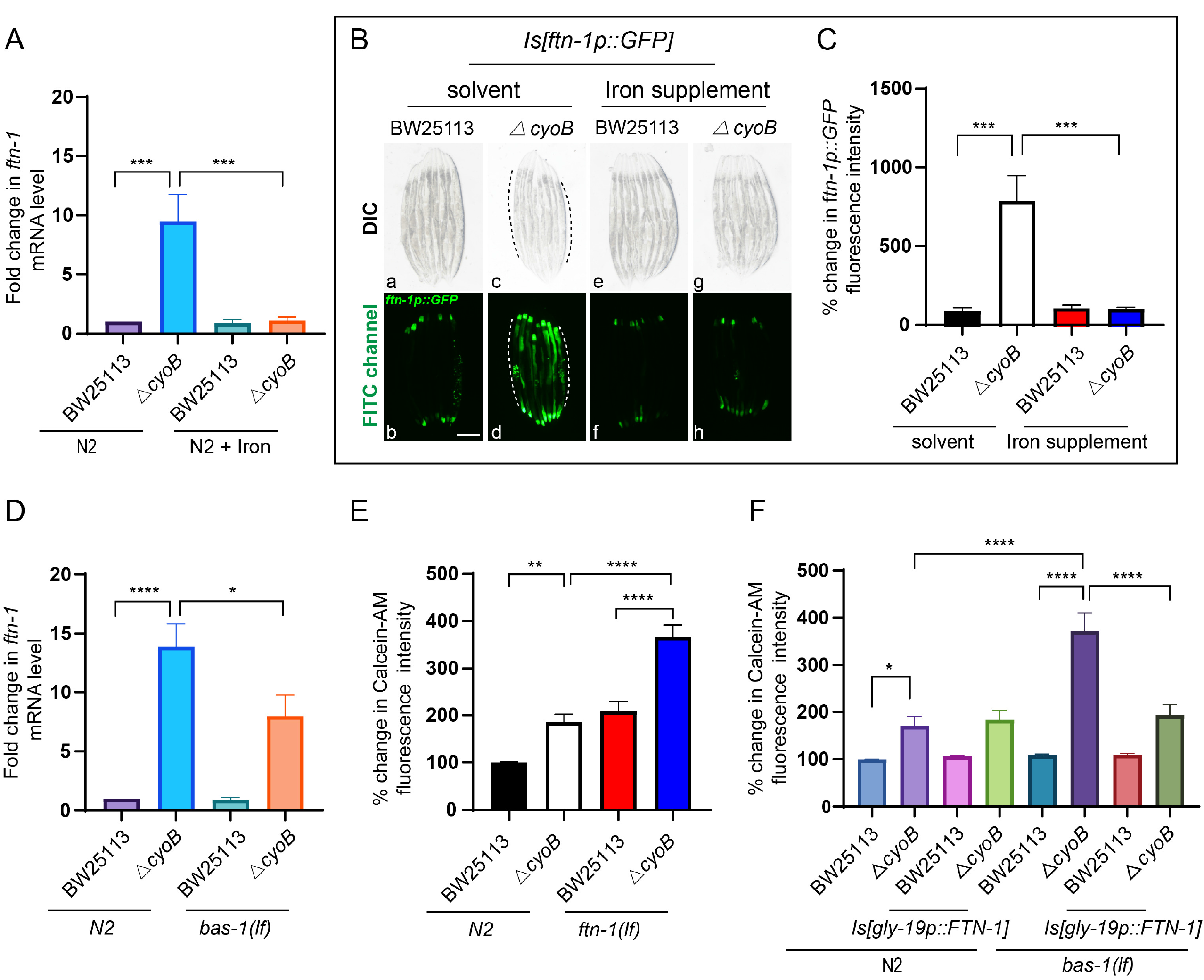
Increased expression of *bas-1* in dopaminergic and serotonergic neurons promotes intestinal *Ferritin (ftn)-1* expression to attenuate the bacteria-induced reduction in labile iron levels. **(A)** qPCR analyses showing that *ftn-1* transcription is increased in response to *ΔcyoB* bacteria, which is reversed by 4mM iron supplementation. The *ftn-1* levels in wild-type N2 worms with the indicated treatments were measured. The fold change was calculated by normalization to the *ftn-1* levels in the wild-type N2 worms treated with BW25113 bacteria. n > 200 each group. (**B and C**) Microscopic images and bar graph illustrating that transcription of *ftn-1* is induced in response to the reduction in labile iron caused by *ΔcyoB E. coli*. An integrated transcriptional fusion reporter expressing *ftn-1p::GFP* was used to indicate *ftn-1* transcription. Dashed lines indicate the intestine. FeCl_3_ supplementation was performed to recover the labile iron level. Fold change in *ftn-1p::GFP* was calculated by normalizing to the levels in animals treated with both BW25113 and solvent. n > 20 for each group. Scale bar, 200 μm. (**D**) qPCR analyses indicating that the increase in *ftn-1* expression in response to bacteria-induced iron reduction is partially dependent on *bas-1*. The fold change was calculated by normalization to the *ftn-1* levels in the wild-type N2 worms treated with BW25113 bacteria. n > 200 each group. (**E**) Bar graph showing that deletion of *ftn-1* exacerbates the reduction in labile iron levels caused by *ΔcyoB* bacteria. Calcein-AM staining was performed to indicate the labile iron level in the worms with the indicated genotype and treatment. The fold change in Calcein fluorescence intensity was analyzed by normalization to the levels in wild-type N2 animals treated with BW25113 bacteria. n > 30 each group. (**F**) Bar graph indicating that recovery of *ftn-1* expression in the intestine is able to suppress the *bas-1(lf)*-caused exacerbated reduction in labile iron in the animals treated with *ΔcyoB* bacteria. The *gly-19* promoter was used to drive the intestinal expression of *ftn-1,* and the Calcein fluorescence intensity was quantified to indicate the labile iron level in *C. elegans* with the indicated treatments and genotypes. The fold change in Calcein fluorescence intensity was analyzed by normalization to the levels in wild-type N2 animals treated with BW25113 bacteria. n > 20 each group. For all panels, experiments were performed for at least three times. Data are the mean ± SEM. *p < 0.05, **p < 0.01, ***p < 0.001 and ****p < 0.0001 by one-way ANOVA.

Furthermore, we investigated the role of this neuron*-*mediated induction of intestinal ferritin expression in modulating iron metabolism. First, we tested whether this increase in *ftn-1* expression in the intestine was responsible for the reduction in labile iron levels observed in the *ΔcyoB E. coli*-treated worms, as ferritin is well known to sequester labile iron for iron storage. Surprisingly, we found that instead of alleviating labile iron insufficiency, *ftn-1(lf)* caused a further reduction in labile iron levels in the animals treated with *ΔcyoB E. coli,* when compared to the *ΔcyoB E. coli*-treated wildtype worms and the *ftn-1(lf)* mutant worms cultured with wildtype bacteria (Fig. 7E). Therefore, these results suggest an unexpected role of the increased *ftn-1* expression in the *ΔcyoB E. coli*-treated worms in attenuating the reduction in labile iron levels, which is consistent with the positive effect of *bas-*1 in promoting *ftn-1* expression to elevate labile iron levels.

Next, to further test the idea that when confronting *ΔcyoB E. coli,* increased *ftn-1* transcription might actually mediate the role of the dopaminergic and serotonergic neurons in elevating the intestinal labile iron level, we investigated whether overexpressing *ftn-1* in the intestine was sufficient to reverse the *bas-1(lf)*-induced exacerbated reduction in labile iron levels in animals treated with *ΔcyoB E. coli*. We constructed an integration transgene that drives the overexpression of *ftn-1* with an intestine-specific promoter, *gly-19*. Indeed, this intestine-specific expression of *ftn-1* was able to suppress about 50% of the increase of Calcein signal resulting from labile iron insufficiency in *bas-1(lf)* worms cultured with *ΔcyoB E. coli* (Fig. 7F), corroborating that intestinal *ftn-1* acts downstream of the dopamine and serotonin signaling to alleviate bacteria-caused decrease of available iron. Moreover, similar to the effect of *bas-1(lf)*, intestinal *atfs-1* knockdown and neuronal *cwn-2* knockdown also inhibited the induction of *ftn-1* by *ΔcyoB E. coli* treatment (Supplementary information, Fig. S7 E-F), further supporting that neuronal signaling triggered by labile iron deficiency and intestinal mitochondrial impairment modulates intestinal *ftn-1* expression. These results together clearly indicate that the increase in *ftn-1* expression in response to *ΔcyoB E. coli* insult acts to promote labile iron level, though we have not been able to illustrate the possible mechanism. In summary, our findings uncover an unexpected gut-brain-gut axis, in which head dopaminergic and serotonergic neurons detect the bacteria-caused insufficiency of labile iron levels through increased *bas-1* expression by perceiving the resultant impairment of the intestinal mitochondria and in turn coordinate the expression of intestinal *ftn-1* to facilitate the elevation of labile iron in the gut (Fig. 8).

**Fig. 8:**
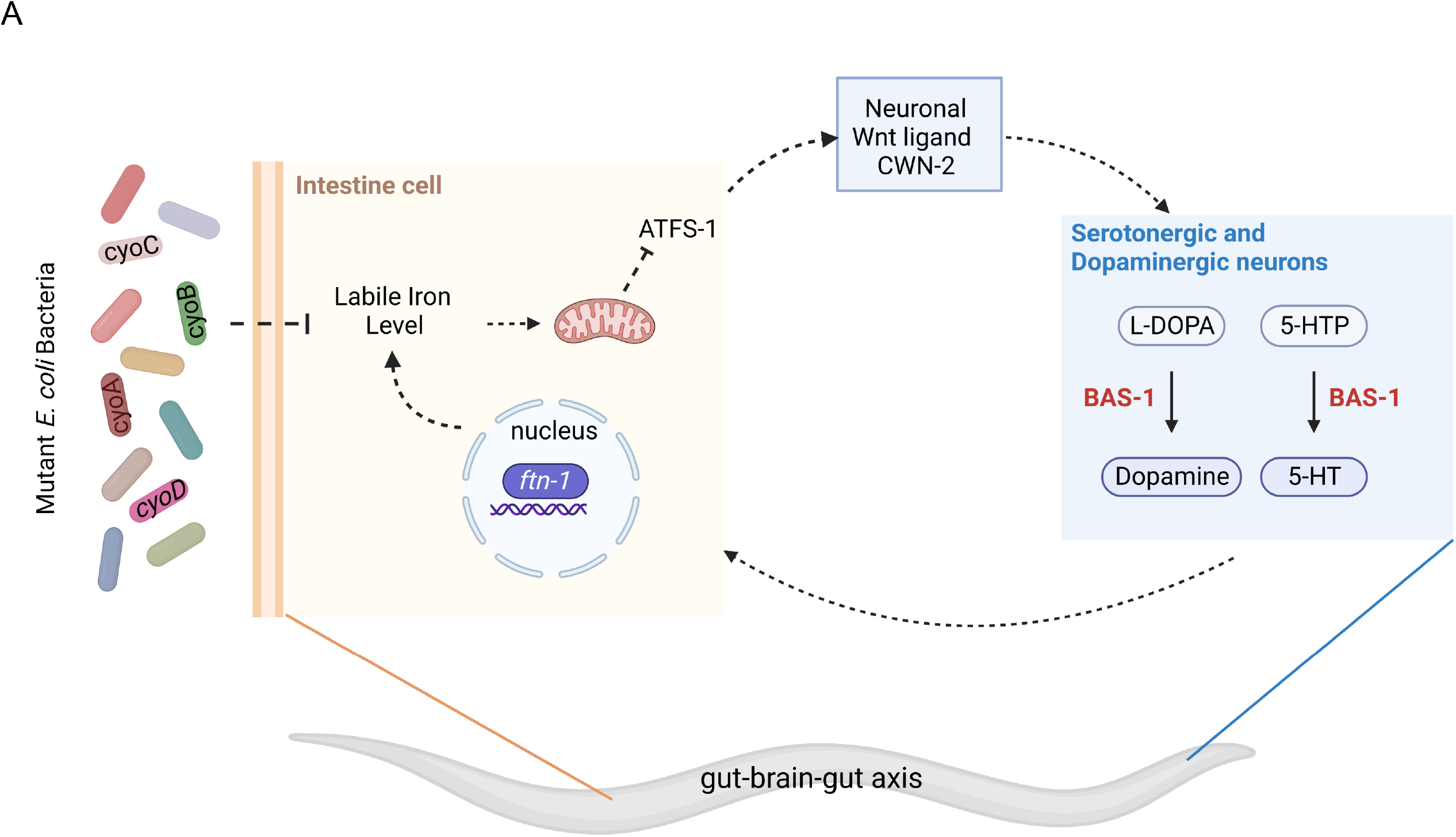
Schematic model for a gut-brain-gut axis mediating the role of host brain in surveilling the bacteria that reduce the labile iron level. **(A)** Dopaminergic and serotonergic neurons are able to perceive bacterial activity changes, such as inactivation of respiratory chain genes, by detecting bacteria-caused reductions in labile iron levels in the intestine of the host to increase dopamine and serotonin levels and then orchestrate the host response to cope with peripheral iron level insufficiency, which is important for preventing further reductions in labile iron levels and mitochondrial function decline. Specifically, dopaminergic and serotonergic neurons respond to the bacteria-induced reduction in labile iron levels by detecting the resultant impairment of the intestinal mitochondria by elevating the dopamine and serotonin biosynthesis via increasing the expression of BAS-1. ATFS-1, a master mitochondrial stress sensor, functions in the gut to transmit this mitochondrial function decline signal from the gut to these BAS-1-expressing neurons, which is at least partially mediated by neuronal CWN-2 Wnt. Subsequently, this intestinal-ATFS-1-mediated induction of BAS-1 expression orchestrates the brain response to cope with the bacteria-caused reduction in labile iron levels by promoting intestinal *ftn-1* expression. Therefore, we identified a gut-brain-gut axis by which the host is able to monitor the bacterial activity change by perceiving the consequent iron metabolism alterations.

## Discussion

What and how alterations of bacterial activities could be detected by the animal brain and the relevant physiologic significance are underexplored. In this study, by using *C. elegans* as a model system, we aimed to systematically understand what changes in the bacterial activity could trigger responses from the head dopaminergic and serotonergic neurons to elevate the biosynthesis of dopamine and serotonin as well as the underlying mechanisms. We screened for single deletions of all *E. coli* nonessential genes that promote the expression of BAS-1, a shared enzyme for dopamine and serotonin biosynthesis, in the head neurons of *C. elegans* and discovered 29 mutations in bacterial genes involved in various processes, including the respiratory chain, that could trigger response from these neurons. Interestingly, we found that these neurons perceive the bacteria with deletions of bacterial respiratory chain genes by inspecting the resultant reduction in labile iron levels in the host. Mechanistically, this bacteria-induced labile iron scarcity caused mitochondrial impairment in the gut to trigger neuronal BAS-1 expression, and this process is critically mediated by the intestine-expressed mitochondrial stress sensor ATFS-1; this neuronal response subsequently coordinates responses to promote intestinal ferritin (FTN-1) expression, which displayed the unexpected effect of elevating the labile iron level. The CWN-2 Wnt originating from neurons mediates the communication from the intestinal cells to dopaminergic and serotonergic neurons, thus representing a novel cell-nonautonomous signal that modulates neurotransmitter biosynthesis and systemic iron homeostasis. Functionally, the responses from the dopaminergic and serotonergic neurons are crucial for animals to maintain iron homeostasis when animals confront bacteria-induced iron insufficiency, and inhibiting dopamine and serotonin signaling causes a further reduction in labile iron levels as well as an exacerbated decline in mitochondrial function. In summary, we reveal that serotonergic and dopaminergic neurons monitor changes in a variety of bacterial processes, providing a comprehensive understanding of what bacterial activity alterations affect the brain. Furthermore, we discovered a gut-brain-gut axis (intestinal ATFS-1-neuronal BAS-1-intestinal FTN-1) (Fig. 8), which not only represents a new regulatory mechanism for monitoring and re-establishing iron homeostasis but also exemplifies a bi-directional process underlying the crosstalk between the host nervous system and bacteria.

Our study provides a fascinating perspective to investigate impacts of changes in bacterial activity on altering the brain activity. We performed a genome-wide screen and revealed single deletions of 29 bacterial genes involved in a variety of processes that provoke a robust response in *C. elegans* serotonergic and dopaminergic neurons. According to our knowledge, our study presents the first example of an overall understanding of the impact of bacterial activity on the nervous system and the positive hits from the screen in Figure S1 should provide an important resource for other researchers in the field to identify bacterial factors that modify the dopaminergic and serotonergic neurons. Furthermore, despite intensive research in the field, the detailed cellular and molecular mechanisms underlying bacteria-brain interactions, including the information needed to be communicated, remained largely elusive. Our study identified a gut-brain-gut axis that mediates the bacteria-host communications and also revealed that the level of intracellular labile iron is an important mediator that can convey changes in bacteria to the host nervous system, thus providing mechanistic insights into the microbiota-brain axis. Since the genes in this axis, such as ferritin in iron metabolism, CWN-2 Wnt and serotonin- and dopamine-related signaling, are highly conserved, it is conceivable that a similar axis may also exist to connect the brain with the gut to modulate iron metabolism as well as to mediate the role of brain in monitoring bacterial activity changes in other animals.

During the interactions with the host, microbes often compete for the essential iron, a growth limiting nutrient in the environment ^21,27,39^; thus, to counteract the bacteria-induced iron loss, mechanisms have evolved in the host to tightly monitor the iron level. In particular, interorgan regulation of iron metabolism that coordinates iron availability of different tissues is crucial for maintaining systemic iron homeostasis under this iron scarcity condition and the liver was the only known organ that is capable of coordinating iron metabolism in different tissues through secreting hepcidin ^32,40^. Here, we discovered a surprising role of the nervous system in modulating iron metabolism through a gut-brain-gut axis and that Wnt acts as a previously-unknown modulator of systemic iron metabolism, which represents a newly identified interorgan communication mechanism for maintaining iron homeostasis at the whole animal level. The head neurons monitor iron insufficiency by inspecting the induced mitochondrial impairment, suggesting that this gut-brain-gut axis is likely to detect relatively severe and chronic reductions in iron levels. The gut is the major organ for absorbing iron as well as a battlefield for competing with bacteria for iron; thus, modulating intestinal iron metabolism by the nervous system holds the capacity to maintain body iron homeostasis. Due to the severe pathological consequences resulted from both insufficient or excessive body iron ^32^, our discovery of the role of the nervous system in monitoring iron metabolism changes in peripheral tissues and mediating the important processes of host-bacteria iron communication is highly important, which deepens our understanding of interorgan regulation of systemic iron homeostasis and the mechanisms underlying the surveillance of the gut microbial content by the nervous system.

Surprisingly, we found that ferritin, well-known to sequester and store iron ^41^, could play an opposite role in elevating free iron levels when worms were treated with the *ΔcyoB E. coli.* Actually, increased ferritin expression has also been detected in humans during infection or under iron-insufficient conditions ^42,43^, and it is worth exploring whether this increased ferritin level may also act to counter further reduction in iron. These observations implicate that the physiologic context, such as the bacteria challenges, may affect what functions ferritin exerts; however, we have not been able to elucidate the mechanism underlying the role of ferritin in increasing labile iron in this study. Maintaining iron homeostasis is crucial to prevent pathogenesis resulting from iron deficiency or overload; thus, it will be highly important to investigate whether and how bacteria may affect the role of ferritin in modulating labile iron levels, which would deepen our understanding of the mechanism by which ferritin functions.

The impact of gut bacteria on the human brain has been suggested to cause severe pathological consequences, such as neurologic defects. The identification of various bacterial processes affecting head neurons in this study supports the idea that when studying the cause of brain disorders, in addition to analyzing the alterations in the brain, changes in the activities of the associated bacteria and the peripheral tissues may also need to be considered as the original contributors for certain neurologic defects ^1^. Furthermore, dopamine and serotonin are important neurotransmitters whose deficiency results in various neurological defects, such as depression and neurodegeneration, and supplementing the neurotransmitter precursors that cross blood-brain barrier is clinically used to treat these disorders. The bacteria strains uncovered in this study to stimulate the biosynthesis of serotonin and dopamine may hold great potential for developing therapeutic strategy for handling neurologic diseases caused by the decrease in serotonin and dopamine. Moreover, our findings may stimulate new studies to investigate how brain defects could cause iron imbalance as well as whether iron insufficiency alters brain activity in other animals. For example, an obvious symptom of iron deficiency anemia in human is the mysterious pica, which is an unusual eating habit or craving for nonnutritive substances ^44^, supporting that iron insufficiency alters brain activity in humans. Our research provides the first mechanistic study showing how iron insufficiency affects head neurons, which may shed light on the connection between iron deficiency and brain disorders.

## Materials and methods

### Strains and worm maintenance

The following bacterial strains were used in this study: *E. coli* OP50 obtained from the Caenorhabditis Genetics Center (CGC), *E. coli* K-12 Keio Knockout library collection (OEC4988), the corresponding wild-type parent strain BW25113 (OEC5042) and the ORF RNAi Collection (RCE1181) were from Dharmacon. The following *C. elegans* strains were used in this study: wild-type N2, LC33 *[bas-1(tm351)]*, CB1111 *[cat-1(e1111)]*, CB1112 *[cat-2(e1112)]*, MT14984 *[tph-1(n4622)]*, VC3201 *[atfs-1(gk3094)]*, QC115 *[atfs-1(et15)]*, RB2603 *[ftn-1(ok3625)]*, GA631 *[lin-15B&lin-15A(n765); wuIs177(ftn-1p::GFP + lin-15(+)]*, SJ4100 *[hsp-6p::GFP + lin-15(+)]*, *MT1215 [egl-20(n585)]*, VP303 *[rde-1(ne219)]; kbIs7 [nhx-2p::rde-1 + rol-6(su1006)]*, TU3401 *[sid-1(pk3321)]; pCFJ90 [(myo-2p::mCherry) + unc-119p::sid-1]*, VC636 *[cwn-2(ok895)]*, MAT137 *jefEx46[gly-19p::ATFS-1^△1–32 myc^]; Is[bas-1p::BAS-1::GFP]*, MAT138 *jefIs8[gly-19p::FTN-1::unc-54 3’UTR]*, MAT140 *jefIs8 [gly-19p::FTN-1::unc-54 3’UTR]; [bas-1(tm351)]*, MAT141 *[rde-1(ne219)]; [nhx-2p::rde-1 + rol-6(su1006)]; Is[bas-1p::BAS-1::GFP]*, MAT154 *jefEx46[gly-19p::ATFS-1^△1–32^ ^myc^]* and MAT193 *[cwn-2(ok895)];Is[bas-1p::BAS-1::GFP].* The *Is[bas-1p::BAS-1::GFP]* strain was obtained from the Shiqing Cai laboratory. MAT137, MAT141, MAT138, MAT140, MAT154 and MAT193 were made in our laboratory. All other strains were from the CGC. All *C. elegans* strains were grown at 20 ℃ on nematode-growth media (NGM) plates seeded with OP50 unless otherwise indicated.

### Screening for the *E. coli* bacterial genes whose single deletion promotes BAS-1::GFP expression in worm neurons and GO analysis

*Bas-1* is specifically expressed in dopaminergic and serotonergic neurons, and the change in BAS-1 expression was analyzed to evaluate the response of these neurons to the presence of bacteria. A BAS-1::GFP translational fusion reporter strain, a gift from Shiqing Cai’s lab ^26^, and the Keio library collection (OEC4988) containing 3985 single deletions of all the nonessential genes of *E. coli* were used to screen for the *E. coli* mutants that increase BAS-1::GFP expression of the host. Totally three rounds of screening were performed to determine the bacterial gene deletions that enhance BAS-1::GFP expression. For each round, each *E. coli* mutants were cultured in 200 μL liquid broth with 100 μg/ml kanamycin overnight using 96-well plates and then seeded on NGM plates containing 50 μg/ml kanamycin and the BAS-1::GFP was acquired with Nikon DS-Qi2 microscopy. As previously indicated ^26^, an increase in BAS-1::GFP by ≥10% relative to animals feeding with parental BW25113 wild-type bacteria was considered to be significant. The 432 bacterial clones identified as increasing BAS-1::GFP expression in the first round were re-examined in the second screening round. Then, the 70 positive clones from the second round were included in the third round and 29 positive clones were observed to enhance BAS-1::GFP expression in all three independent experiments. To explore what bacterial processes would affect host BAS-1::GFP expression, GO analyses for these 29 mutant genes were then performed by clusterProfiler, and corresponding images were generated by ggplot2 (v3.3.0).

### Imaging and analysis of the expression of transgenic reporters

For microscopic analyses, approximately 150 L1-staged worms expressing indicated transgenic reporters were cultured with indicated *E. coli* bacteria, and when the worms reached the day-1 adult stage, worms were manually picked and anesthetized with 150 mg/mL levamisole on a 2% agarose pad before imaging. For sample randomization, the worms analyzed by complex microscopy were randomly selected under a dissection microscope. Fluorescence levels in the head neurons of BAS-1::GFP-expressing worms in all the experiments, except the ones from the screen in Figure 1 B-C and Figure S1 that were scored by using Nikon DS-Qi2 microscopy, were acquired by confocal microscopy (Zeiss LSM 880 NLO, 40X) and ImageJ software was used to quantify the fluorescence intensity. To analyze the expression of *ftn-1p::GFP* and *hsp-6p::GFP* resulting from specific bacterial treatments, pick and anesthetize worms on NGM plate with 8-10 animals per group to image with a Nikon SMZ18 or mount worms on slides for imaging using a Nikon DS-Qi2 microscope. The intensity of GFP fluorescence signal was quantified with ImageJ.

### Analyzing levels of dopamine and serotonin by HPLC-MS

The dopamine and serotonin levels in *C. elegans* were detected by using HPLC-MS. Approximately 2,000-10,000 day-1 adult worms treated with BW25113 or *ΔcyoB E. coli* were collected and washed with M9 buffer. Add 80% HPLC-grade methanol (pre-chilled at −80°C) to the pelleted worms with the volume/weight ratio at 10 μL/mg and then homogenize the worms with FastPrep-24 (MP Biomedicals) for 30 seconds. Next, transfer the worm lysates to a new eppendorf tube and incubate at −80°C overnight to allow extract the dopamine and serotonin. Then, centrifuge these samples at 14,000 g for 20 min at 4°C and transfer the supernatant that contains the dopamine and serotonin components to a new eppendorf tube. 10 μL of these supernatants were used to determine the protein level using the BCA Protein Assay Kit (E162-01). The rest of the supernatants were lyophilized in a vacuum centrifugal concentrator (MS-SP102) and the dried powder was resuspended in 60 μL of 80% methanol followed by vigorous vortex. After centrifugation at 14,000 g, 10 μL of the resulting supernatant was analyzed by HPLC–MS to determine the dopamine and serotonin levels. A standard curve of the commercially obtained standards (Sigma, USA) was used to determine the dopamine and serotonin concentrations.

### Chemical supplementation

FeCl_3_ (Sigma‒Aldrich Cat# 157740), dopamine (Sigma‒Aldrich Cat# H8502) and serotonin creatine sulfate (Sigma‒Aldrich Cat# H9523) dissolved in water and 2-2’ bipyridyl (BP) (Sigma‒Aldrich Cat# D216305) dissolved in DMSO were added to the NGM before pouring to final concentrations of 4 mmol/L, 12.5 mg/ml, 10 mg/ml and 50 μM, respectively. An equivalent volume of the corresponding solvent was added to the NGM as a negative control. To assay the impact of these chemicals on the indicated phenotypes, L1-staged worms were cultured on NGM plates with the indicated chemical supplements and bacterial treatments, and when they reached the day-1 adult stage, the indicated phenotypes were scored as described.

### Labile iron detection by Calcein-AM

The level of intracellular labile iron in *C. elegans* was analyzed with the widely used Calcein-AM dye (Thermo Fisher, Cat #C3100MP) as previously described ^21^. Approximately 20-30 day-1 adult worms with the indicated genotype and bacterial treatment were manually picked under a dissection microscope and allowed to crawl on unseeded NGM plates for one hour to remove the residual bacteria. Then, these worms were transferred to 100 µL of the commonly used M9 buffer (3.0 g KH_2_PO_4_, 6.0 g Na_2_HPO_4_, 0.5 g NaCl, 1.0 g NH_4_Cl dissolved in 1 L H_2_O) containing 0.5 µg/mL Calcein-AM and incubated for two hours at 20℃. Next, these worms stained with Calcein-AM were rinsed with M9 buffer to remove the residual Calcein-AM dye. After the wash, the worms were mounted on an agarose pad (2% agarose), and the Calcein signal was detected under a fluorescence microscope (Nikon DS-Qi2, 40X) with the DIC and FITC channel. The intensity of Calcein fluorescence was analyzed by using ImageJ software.

### Measurement of basal oxygen consumption rate (OCR)

The basal OCR of worms was analyzed by a Seahorse XFe96 analyzer with extracellular flux assay kit as previously described ^45^. Specifically, 200 adult worms with the indicated genotypes and treatments were collected for each group after the residual bacteria were removed by washing with M9 buffer twice. Then aliquot 10-30 worms/well to a seahorse XF96 cell culture microplate. Approximately 5-8 technical replicates per independent experiment and at least three biological replicates were performed. The basal OCR in live *C. elegans* was probed using the following program: 5 cycles of mix for 2 min, wait for 0.5 min, measure for 2 min. The seahorse measurements of OCR were normalized by worm numbers.

### RNA interference in *C. elegans*

RNAi experiments were carried out by feeding the worms with bacteria that generate dsRNA against corresponding worm genes as described previously ^46^. Specifically, the indicated RNAi HT115 *E. coli* bacteria from the MRC RNAi library (Source BioScience) and ORF RNAi library (GE Dharmacon) were cultured in LB medium containing 100 μg/ml ampicillin and 15 μg/ml tetracycline for 8-10 hours at 37°C and then 350 μL RNAi bacteria were seeded onto NGM plates with 2 mM IPTG. Then, these plates with the indicated RNAi bacteria were maintained at room temperature for 3-5 days to allow the production and accumulation of dsRNA before the worms were seeded. Approximately 100-200 L1-staged worms with the indicated genotypes and treatments were cultured on these plates, and the day-1 adults were scored.

HT115 bacteria containing the empty PL4440 vector were used as the RNAi negative control. The VP303 worm strain, which expressed *rde-1* only in the intestine ^34^, was used to perform intestine-specific RNAi analyses. While the TU3401 *C. elegans* strain, which overexpressed *sid-1* by neuron specific promotor in the *sid-1(lf)* mutant, displaying neuron-specific RNAi efficacy ^47^, was used in experiments that aimed to knock down genes specifically in the neurons.

### Analyses of worm survival rate after paraquat treatment

The rate of worms surviving paraquat treatment was analyzed to evaluate the animals’ ability to cope with mitochondrial stress as previously described ^35^. Briefly, L1-staged worms with the indicated genotypes were grown on NGM plates containing BW25113 *or Δ cyoB E. coli* with and without FeCl_3_ supplementation (4mM) at 20°C. When they reached the late L4 stage, these worms were transferred to NGM plates supplemented with 5mM paraquat (Sigma) containing OP50 bacteria. The survival of animals on paraquat plates was scored daily until all worms were dead, which was evaluated by the animals showing no response to a light touch with a platinum wire. If worms died of unnatural causes such as desiccation on the side of the dish, they were excluded from the scoring. The mean survival time and the statistics were analyzed by GraphPad Prism. Mean survival time was calculated as the sum of the survival time of each worm with the indicated genotype and treatment/total number of worms of the corresponding genotype and treatment.

### Generation of transgenic strains

The *Is[gly-19p::FTN-1::unc-54 3’UTR]* and *Ex[gly-19p::ATFS-1^△1–32myc^]* transgenes specifically expressing *ftn-1 and atfs-1(gf)* in the intestine were used to evaluate the role of ferritin in alleviating free iron reduction and the impact of the mitochondrial state on modulating *bas-1* expression and free iron levels, respectively. The promoter of *gly-19* was used to drive the expression of these genes specifically in the intestine ^33^. The 5’TTTTTTTCAGTACATTTTTTCATTTCGGGTTCA3’ and 5’CTGGAAATTTAAATTTAATTCTTTGGAATTTAGT3’ primers were used to generate the promoter of *gly-19* by PCR, and *ftn-1* cDNA was amplified by using the 5’ATGTCTCTAGCTCGTCAAAACTATC3’ and 5’TTAATCAGAAAATTCCT-CTTTGTCGAAC3’ primers. The plasmid used for intestine-expression of *atfs-1(gf)* was obtained from Ye Tian’s lab. A DNA mixture containing 20 ng/μL *myo-2p::mCherry* co-injection marker and 50 ng/μL plasmid with intestine-specific expression of the indicated genes were microinjected to obtain the extrachromosomal transgenes. The worms carrying *gly-19p::FTN-1::unc-54 3’UTR* extrachromosomal array were used for construction of *Is[gly-19p::FTN-1::unc-54 3’UTR]* integration transgenes by using UV irradiation method.

### Quantitative RT‒PCR (qPCR) analyses

To perform qPCR measurements of gene transcription, more than 150 young adult worms with the indicated genotypes and treatments were collected for RNA extraction by using a MicroElute Total RNA kit. Approximately 300-800 ng extracted RNA was treated with One-Step gDNA Remover (Transgene) to remove potential genomic DNA contamination and then used to produce complementary DNA by reverse-transcription with the cDNA Synthesis SuperMix kit (Transgene). The generated cDNA was used as the template for qPCR analysis by utilizing Universal qPCR Master Mix (BioLabs). At least three independent experiments were performed to indicate the change in gene transcription. The transcription of a housekeeping gene, *rpl-26*, was used as an internal control as previously described ^46^. The primers for real-time PCR were as follows:

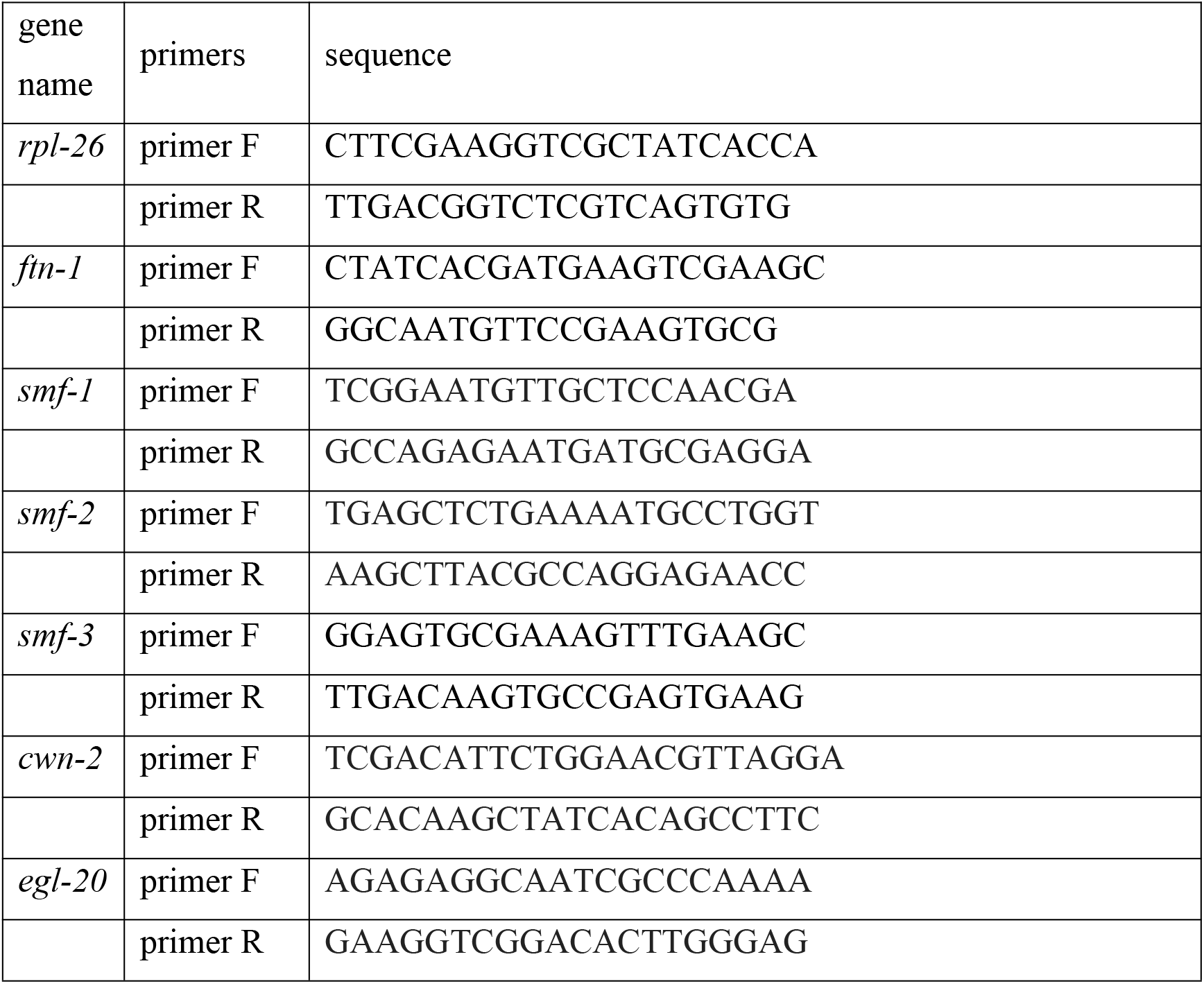

### Statistical analysis

To avoid potential bias, the information on treatment and genotypes was masked before the phenotypes were scored. At least 3 independent experiments were performed for all the results. The sample size (“n” number) indicated in the Figures and Results is the number of worms scored in that experiment. All statistics were analyzed by using GraphPad Prism. Statistical comparisons between two groups and the comparisons among multiple groups were analyzed by Student’s t test and one-way ANOVA, respectively. Data are presented as the standard error of the mean. The significance is indicated as follows: *, p < 0.05, **, p < 0.01, ***, p < 0.001, ****, p < 0.0001.

## Acknowledgments

We thank the CGC (funded by NIH [P40OD010440]) for *C. elegans* strains, S. Cai (Institute of Neuroscience, Chinese Academy of Sciences), Y. Liu (Peking University) and Y. Tian (Institute of Genetics and Developmental Biology, Chinese Academy of Sciences) for providing *C. elegans* strains and reagents, and P. Zhao (Westlake University), S. Wu (Westlake University), F. Jin (Westlake University), the High-performance Computing Center of Westlake University and the Microscopic Imaging Center of Westlake University for technical support. We thank X. Xu and Mass Spectrometry & Metabolomics Core Facility of Westlake University for HPLC-MS analysis. This research was supported by the National Key Research and Development Program of China (No. 2019YFA0802900), National Natural Science Foundation of China (No. 32070565 and No. 31871465), Zhejiang Natural Science Foundation (No. LQ23C040002), HRHI program (No. 202109007 and No. 202209003) of Westlake Laboratory of Life Sciences and Biomedicine and the Westlake Education Foundation.

## Author contributions

H.T. and G.L. conceptualized the study. G.L., Y.W. and X.H. conceived and designed the study. G.L. and Y.W. performed most of the experiments and analyzed the data. X.H. and M.D. carried out part of the experiments and data analyses. H.T., G.L., Y.W. and X.H. wrote the paper. H.T. supervised the study.

---

**Fig. S1:**
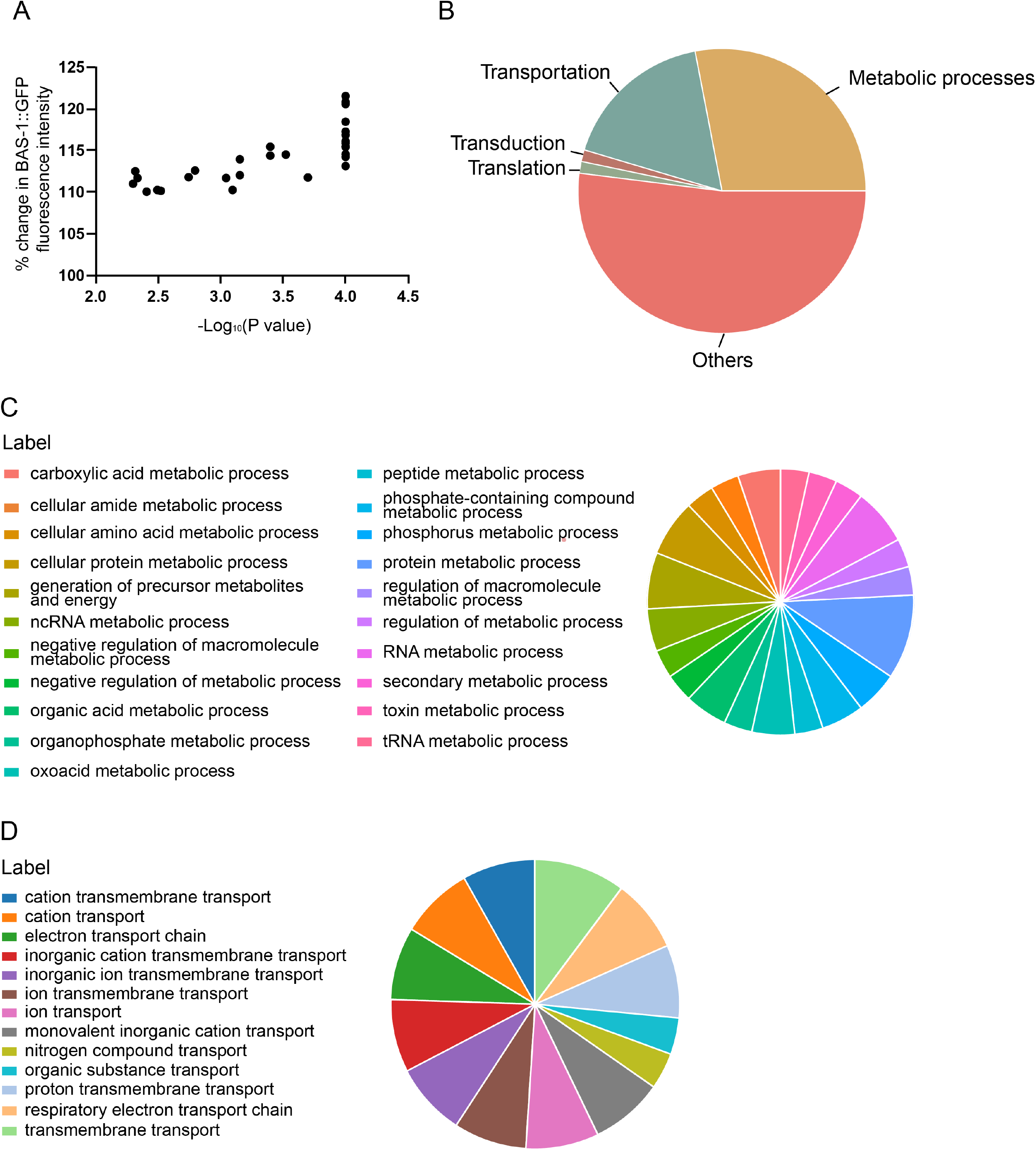
Effects and categories of the bacterial genes that promote BAS-1 expression in *C. elegans* dopaminergic and serotonergic neurons following single deletion. **(A)** Graph showing the increase of BAS-1::GFP induced by the 29 *E. coli* mutants positive hits obtained from the screen described in Figure 1B. % change in BAS-1::GFP fluorescence intensity were normalized to the expression levels in worms treated with wild-type BW25113 *E. coli*. Each dot indicates the effect of one mutant *E. coli* on the BAS-1::GFP level. P values were determined by unpaired two-tailed t test. **(B-D)** Gene ontology analysis showing the enrichment of the screen hits in a variety of bacterial processes, including transportation and metabolism. GO analyses indicate the categories of the positive hits (B). Further GO analysis shows the detailed metabolic processes (C) and transportation processes (D) in which the bacterial genes, whose inactivation promotes BAS-1::GFP expression, are involved.

**Fig. S2:**
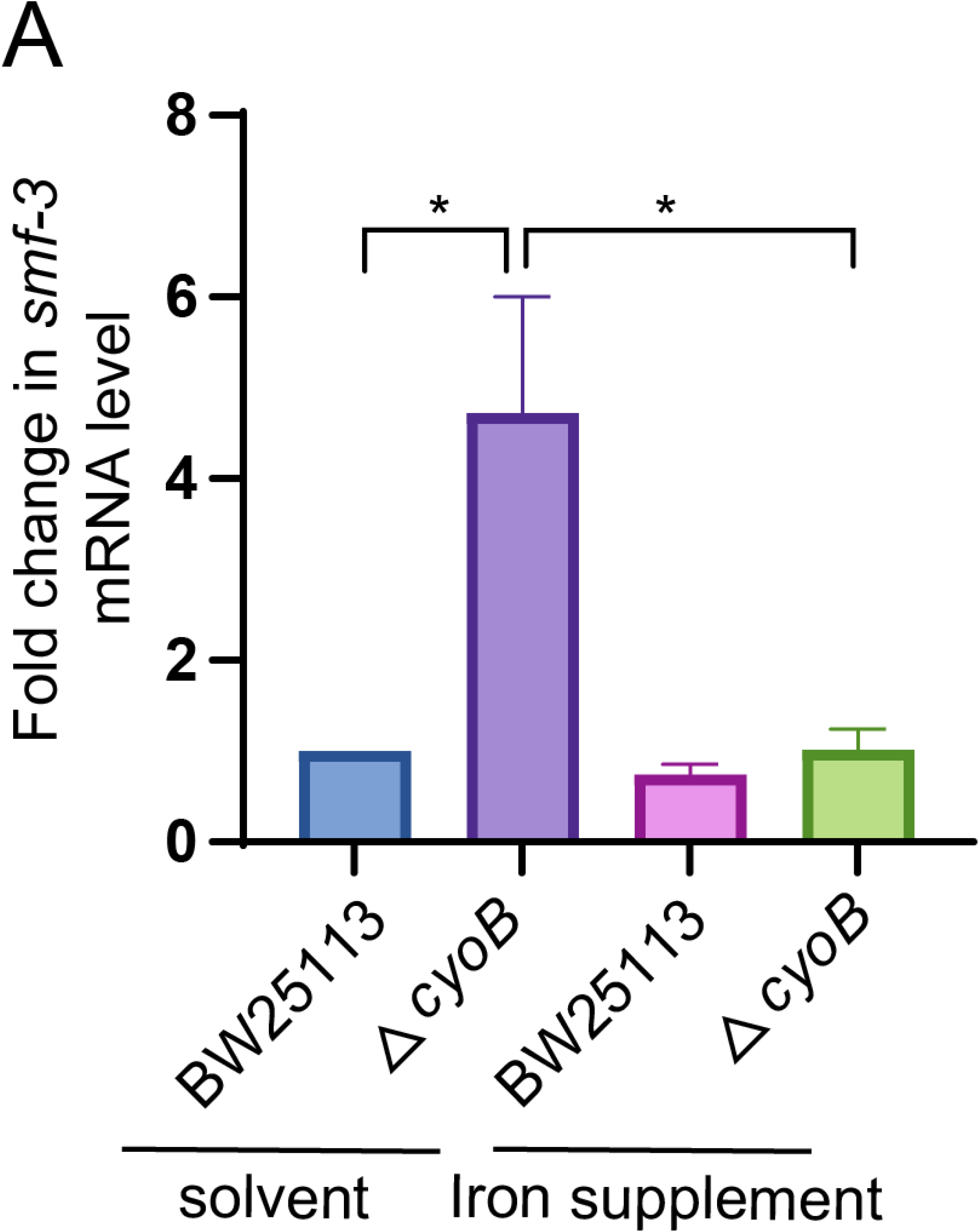
The *ΔcyoB E. coli* treatment induces *smf-3* transcription, which is suppressed by iron supplementation. **(A)** qPCR results showing that transcription of *smf-3*, encoding transporters for iron absorbing in *C. elegans* ^30^, is increased in response to *ΔcyoB E. coli* treatment. 4mM FeCl_3_ was used for iron supplementation. Wildtype animals treated with BW25113 *E. coli* were used as the control to calculate the fold change in *smf-3* mRNA levels. Data are the mean ± SEM. *p < 0.05 by one-way ANOVA; n > 200 each group.

**Fig. S3:**
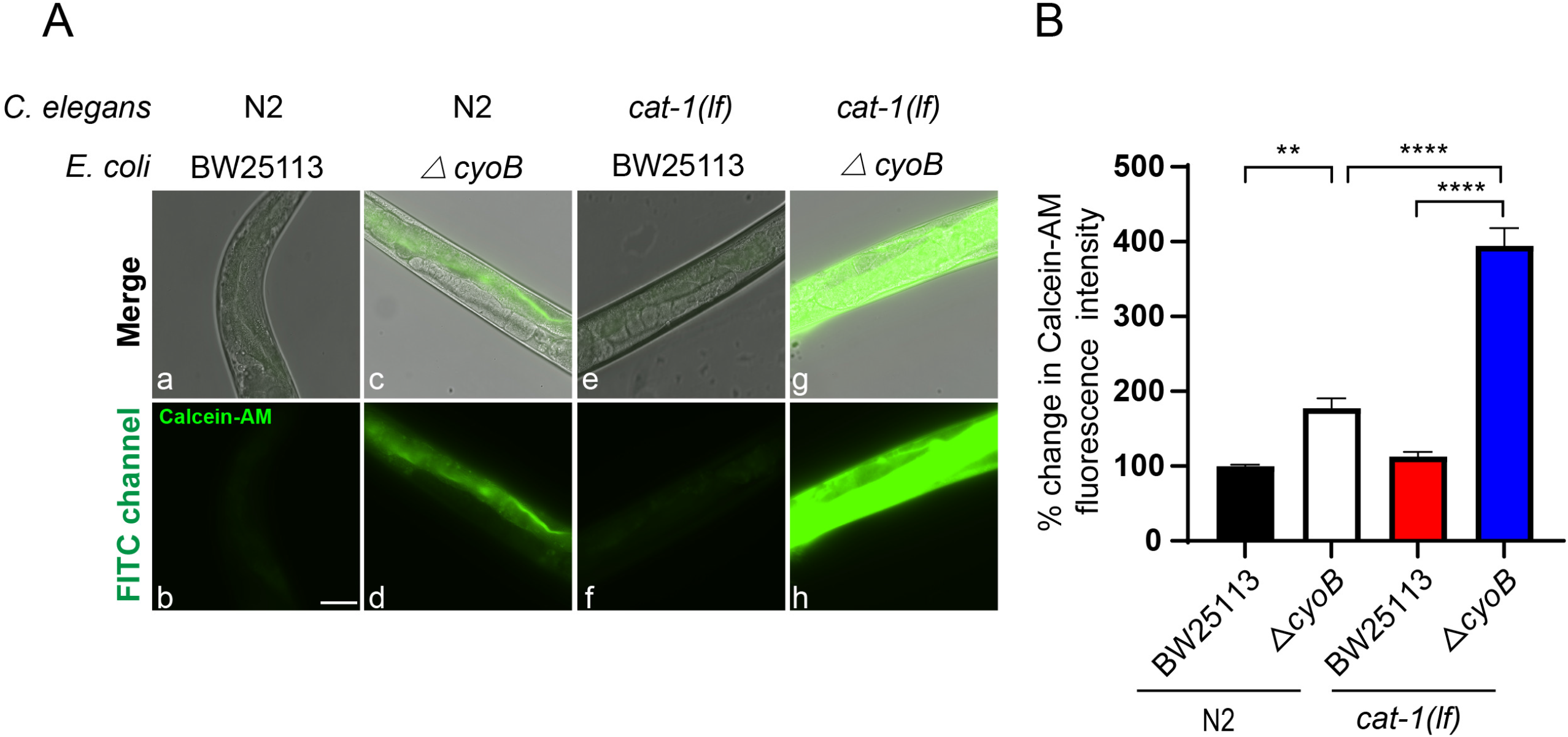
Inhibiting dopamine and serotonin release by blocking their import into vesicles exacerbates the reduction in labile iron caused by *ΔcyoB E. coli*. (**A and B**) Images and bar graph indicating that the *cat-1(lf)* mutation caused a further reduction in the labile iron level in the presence of *ΔcyoB*. The % change in the Calcein fluorescence was calculated by normalization to the levels in wild-type N2 animals treated with BW25113 bacteria. CAT-1 is required for transporting dopamine and serotonin into vesicles for release, and thus, deletion of *cat-1* blocks dopamine and serotonin signaling. Data are the mean ± SEM. **p < 0. 01, ****p < 0.0001 by one-way ANOVA; n > 25 each group. Scale bar, 50 μm.

**Fig. S4:**
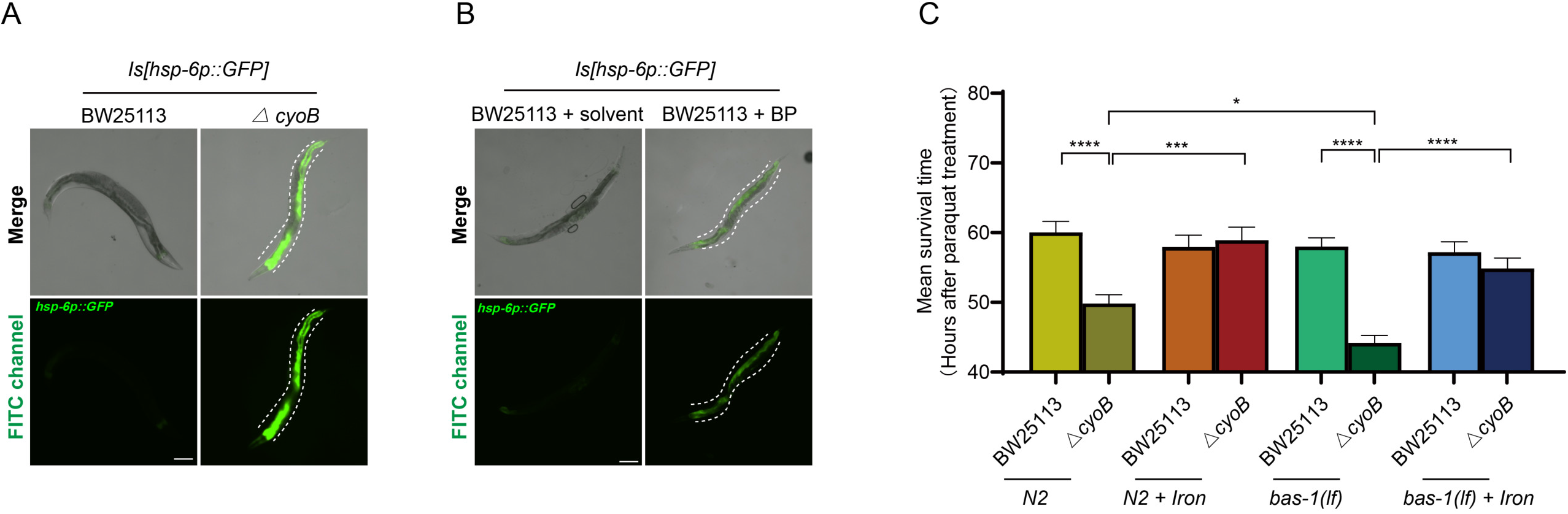
The intestinal mitochondria impairment induced by the *ΔcyoB E. coli* and iron chelator treatment and the role of serotonergic and dopaminergic neurons response to the presence of *ΔcyoB E. coli* in preventing decline in mitochondrial function. (**A**) Microscopic images indicating that *ΔcyoB* mutant *E. coli* mainly induces mitochondrial stress in the intestine of *C. elegans*. The induction of *hsp-6p::gfp* expression in the intestine (white lines) was obviously detected in the presence of *ΔcyoB E. coli*. Scale bar, 100 μm. (**B**) Images showing that the iron chelator BP (25 μM) induces *hsp-6p::gfp* expression mainly in the intestine of *C. elegans*. White lines indicate the location of the intestine. Scale bar, 100 μm. (**C**) Bar graph showing that the increase in *bas-1* expression in response to *ΔcyoB* bacteria is important for preventing a further decline in mitochondrial function. Paraquat was used to challenge mitochondrial function, and the mean survival time of *C. elegans* with the indicated genotypes and treatments was calculated as described in the Methods. Data are the mean ± SEM. *p < 0.05, ***p < 0.001 and ****p < 0.0001 by one-way ANOVA; n > 120 each group.

**Fig. S5:**
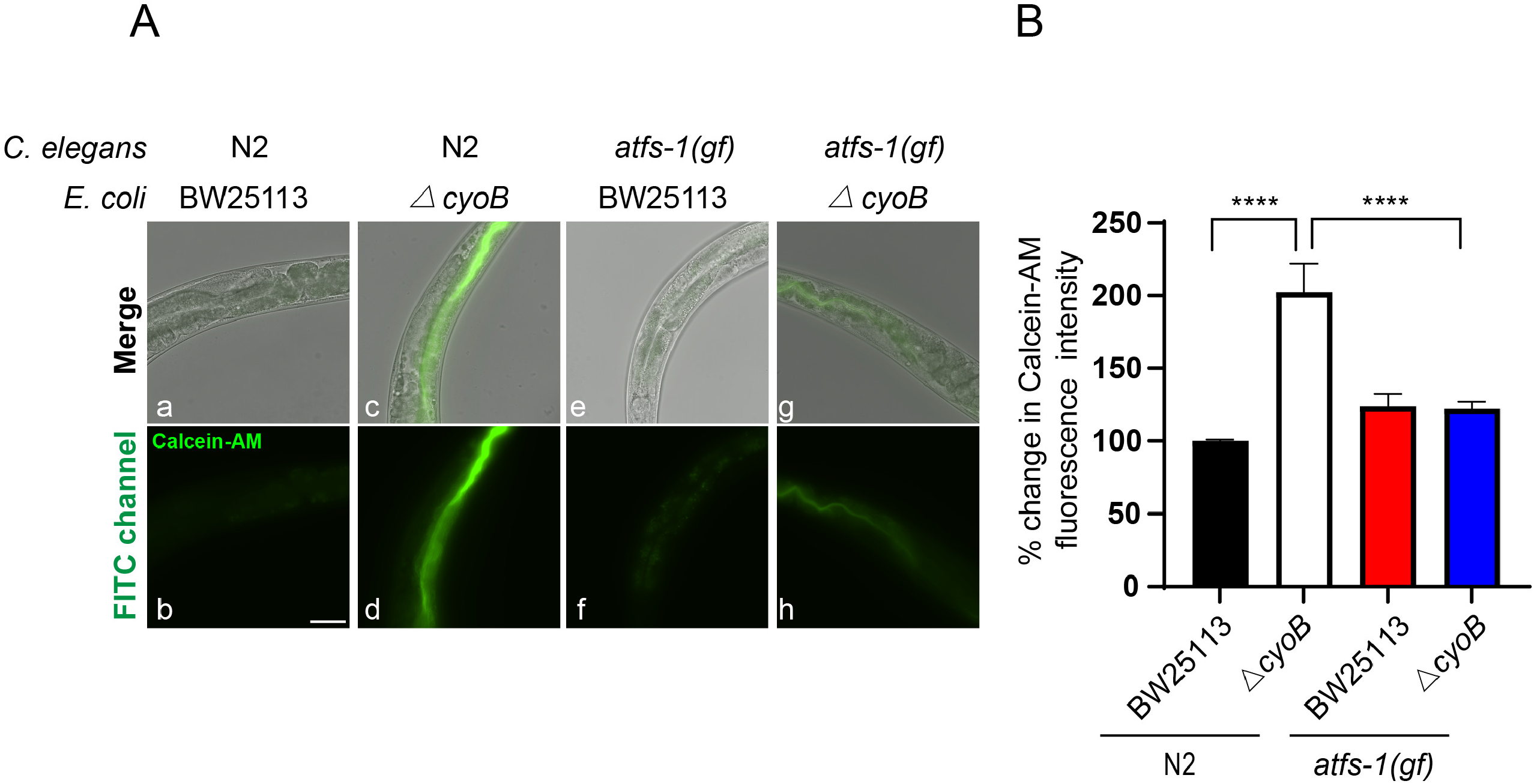
Over-activation of ATFS-1 is able to alleviate the reduction in labile iron levels caused by *ΔcyoB E. coli*. (**A and B**) Images and bar graph indicating that the dominant active form of ATFS-1 suppressed the reduction in labile iron level caused by the *ΔcyoB E. coli* treatment. Calcein-AM staining was performed in N2 and *atfs-1(gf)* worms with the indicated bacterial treatments to indicate the labile iron levels. The % change in Calcein fluorescence was calculated by normalization to the levels in N2 worms treated with BW25113. Data are the mean ± SEM. ****p < 0.0001 by one-way ANOVA. n > 25 each group. Scale bar, 50 μm.

**Fig. S6:**
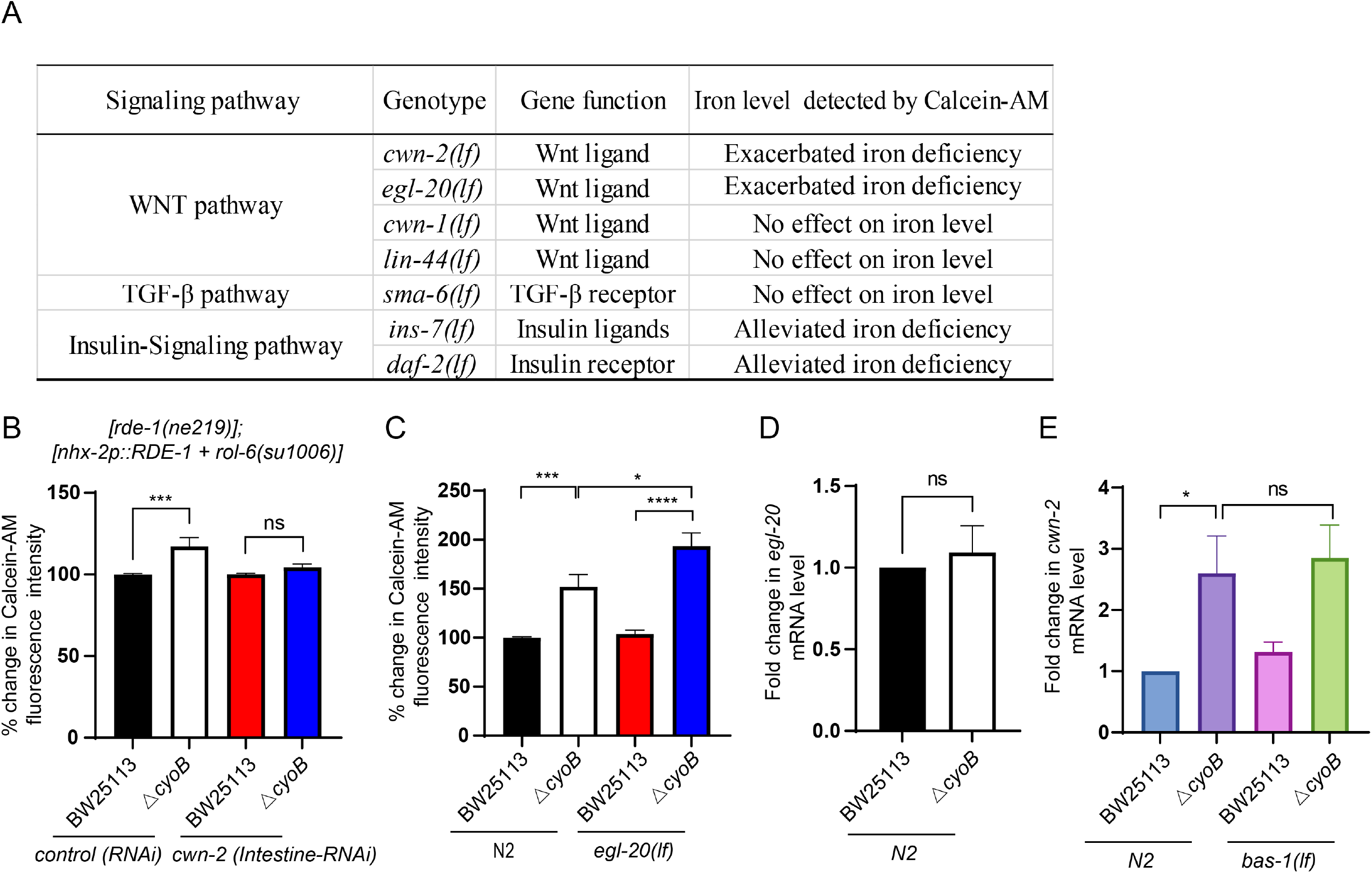
Wnt ligands are required for alleviating iron deficiency caused by the *ΔcyoB E. coli* mutant. **(A)** Table showing the effects of genes in the indicated signaling pathway on bacteria-caused reductions in labile iron levels. Wild-type N2 worms and the listed mutant *C. elegans* were treated with BW25113 and the *ΔcyoB E. coli* mutant, and Calcein-AM staining was then performed to determine the level of labile iron. Exacerbated iron deficiency indicates that the mutant worms treated with *ΔcyoB E. coli* show even lower labile iron levels than both the N2 worms cultured with *ΔcyoB E. coli* and the mutant worms treated with BW25113. There was no effect on the iron level when no obvious difference in the labile iron level between the mutant worms and the wild-type N2 worms after treatment with *ΔcyoB* was observed. When lower Calcein fluorescence was observed in the mutant worms than in the N2 worms after *ΔcyoB* treatment, the effect was indicated to be “alleviated iron deficiency”. **(B)** Bar graph showing that *cwn-2* does not function in the intestine to alleviate the reduction in labile iron levels in the presence of *ΔcyoB*. The VP303 strain, in which RNAi is only effective in the intestine, was used to perform *cwn-2 (intestine-RNAi)* analyses in the presence of BW25113 or *ΔcyoB,* and the Calcein fluorescence intensity was analyzed. The % change in Calcein fluorescence was calculated by normalization to the levels in animals treated with control RNAi and BW25113. Data are the mean ± SEM. ns, p > 0.05 and ***p < 0.001 by one-way ANOVA. n > 20 each group. **(C)** Bar graph displaying that deletion of Wnt ligand *egl-20* causes aggravated reduction in labile iron levels in the presence of *ΔcyoB E. coli*. Worms with the indicated genotypes and treatments were stained with Calcein-AM to analyze the labile iron level. The fold change was calculated by normalizing to the levels in N2 worms treated with BW25113. In contrast to *egl-20(lf) C. elegans* treated with BW25113 or the N2 worms treated with *ΔcyoB,* Calcein-AM analysis showed that *egl-20(lf)* worms treated with *ΔcyoB* leads to an exacerbated reduction in labile iron levels, as evidenced by the increase in Calcein fluorescence. *p < 0.05, ***p < 0.001 and ****p < 0.0001 by one-way ANOVA. n > 30 each group. **(D)** qPCR results show that *egl-20* mRNA levels are not significantly changed in the presence of *ΔcyoB E. coli.* N2 animals treated with BW25113 was used as the control to calculate the fold change in mRNA levels. Data are the mean ± SEM. ns, p > 0.05 by unpaired t test. n > 200 each group. **(E)** qPCR analyses indicate that deletion of *bas-1* shows no effect on the transcription of *cwn-2,* supporting that *bas-1* functions downstream of *cwn-2*. The mRNA of *cwn-2* in worms with the indicated genotypes and treatments was analyzed by qPCR. In the presence of *ΔcyoB E. coli* mutant bacteria, the *bas-1(lf)* worms displayed similar levels of *cwn-2* mRNA to that in wild-type N2 *C. elegans.* N2 wildtype animals treated with BW25113 was used as the control to calculate the fold change in mRNA levels. ns, p > 0.05 and *p < 0.05 by one-way ANOVA. n > 200 each group.

**Fig. S7:**
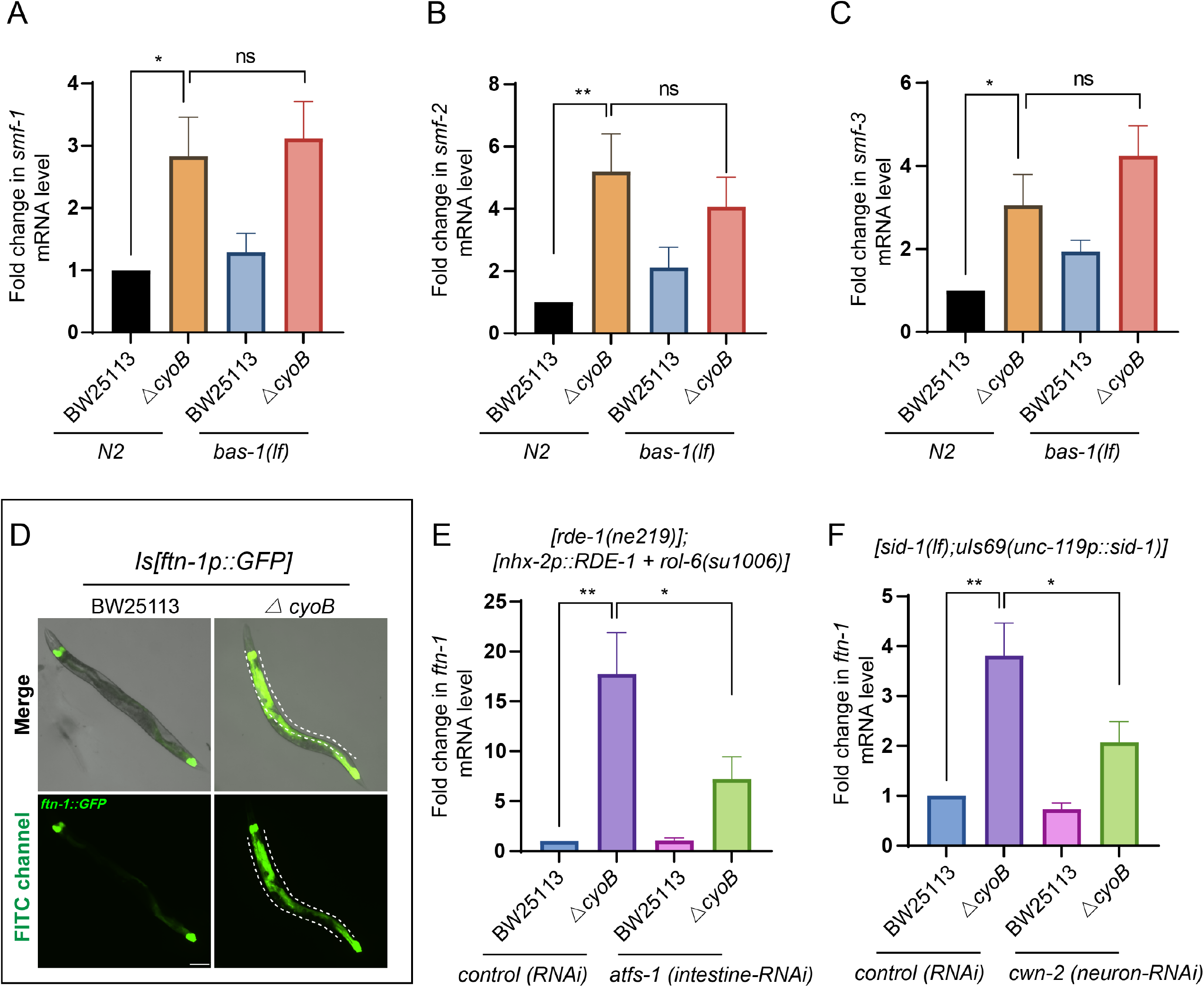
In response to the *ΔcyoB E. coli* mutant, increased transcription of ferritin but not iron absorption genes is dependent on the host neuronal response. **(A-C)** qPCR results show that *bas-1* is not required for the induction of *smf-1/-2/-3* mRNA expression caused by the *ΔcyoB E. coli*. The transcription of *smf-1/-2/-3*, genes encoding transporters for absorbing iron in *C. elegans* ^30^, are increased in response to *ΔcyoB E. coli.* In the presence of *ΔcyoB E. coli,* the *bas-1(lf)* worms showed no significant difference in the level of *smf-1/-2/-3* mRNA compared with N2 wild-type *C. elegans*. Wildtype animals treated with BW25113 was used as the control to calculate the fold change in mRNA levels. n > 200 each group. **(D)** Microscopic images showing the intestine-specific expression of *ftn-1* and its induction by the presence of *ΔcyoB*. The induction of the Is[*ftn-1p::GFP*] was observed with *ΔcyoB* treatment when compared to the BW25113 group. The intestine was outlined. Scale bar, 100 μm. **(E)** qPCR results show that intestine *atfs-1* is at least partially required for the increase in *ftn-1* transcription in response to *ΔcyoB E. coli* mutant bacteria. The VP303 worm strain was used to perform intestine-specific knockdown of *atfs-1* in the presence of the indicated bacteria. Fold change was determined by normalization to the animals fed with both control RNAi and BW25113 bacteria. n > 200 each group. **(F)** qPCR analyses show that neuronal *cwn-2* is at least partially required for the induction of *ftn-1* mRNA expression in the presence of *ΔcyoB E. coli* mutant bacteria. TU3401[*sid-1(lf); uIs69(unc-119p::sid-1)*] was used to knock down *cwn-2* specifically in neurons. *ΔcyoB E. coli*-induced *ftn-1* transcription is partially suppressed by *cwn-2(neuron-RNAi)*. n > 200 each group. For all the panels, data are the mean ± SEM. ns, p > 0.05, *p < 0.05 and **p < 0.01 by one-way ANOVA.

